# Multiplexing cortical brain organoids for the longitudinal dissection of developmental traits at single cell resolution

**DOI:** 10.1101/2023.08.21.553507

**Authors:** Nicolò Caporale, Davide Castaldi, Marco Tullio Rigoli, Cristina Cheroni, Sebastiano Trattaro, Alessia Valenti, Matteo Bonfanti, Sarah Stucchi, Alejandro Lopez Tobon, Dario Ricca, Manuel Lessi, Martina Pezzali, Alessandro Vitriolo, Katharina T. Schmid, Matthias Heinig, Fabian J. Theis, Carlo Emanuele Villa, Giuseppe Testa

**Affiliations:** Department of Oncology and Hemato-Oncology, University of Milan, Via Santa Sofia 9, 20122, Milan, Italy; Human Technopole, Viale Rita Levi-Montalcini 1, 20157, Milan, Italy; Institute of Computational Biology, Helmholtz Zentrum München – German Research Center for Environmental Health, Neuherberg, Germany; Department of Mathematics, Technical University Munich, Munich, Germany

**Author notes:** These authors contributed equally, are listed in alphabetical order, and have the right to list their name first in their CV. These authors jointly supervised this work (Nicolò Caporale, Carlo Emanuele Villa, Giuseppe Testa). PhD students within the European School of Molecular Medicine (SEMM).

## Abstract

The combination of brain organoid and single cell omic technologies holds transformative potential to dissect human neurobiology at high resolution and with mechanistic precision. Delivering this promise in the context of human neurodiversity, physiological and pathological alike, requires however a major leap in scalability, given the need for experimental designs that include multiple individuals and, prospectively, population cohorts. To lay the foundation for this, we implemented and benchmarked complementary strategies to multiplex brain organoids. Following an extended longitudinal design with a uniquely informative set of timepoints, we pooled cells from different induced pluripotent stem cell lines either during organoids generation (upstream multiplexing in mosaic models) or before single cell-RNAseq library preparation (downstream multiplexing). We developed a new method, SCanSNP, and an aggregated call to deconvolve organoids cell identities, overcoming current criticalities in doublets prediction and low quality cells identification and improving accuracy over state of the art algorithms. Integrating single cell transcriptomes and analysing cell types across neurodevelopmental stages and multiplexing modalities, we validated the feasibility of both multiplexing methods in charting neurodevelopmental trajectories at high resolution, linking their specificity to genetic variation between individual lines. Together, this multiplexing suite of experimental and computational methods provides an enabling resource for disease modelling at scale and paves the way towards an *in vitro* epidemiology paradigm.

## Introduction

The polygenic underpinnings of human neurodiversity, in its physiological and pathological unfolding alike, have been eloquently referred to as terra incognita, calling for new maps to trace that unfolding in the authenticity of human genetic backgrounds and thereby render it mechanistically actionable. Developmental stochasticity and environmental triggers add to such complexity, and the increasingly broader range of exposome that is becoming measurable promises to make gene environment interactions finally tractable at meaningful scales ^1–4^.

Towards these overarching goals, brain organoid and single cell multi-omic technologies have afforded major strides in the mechanistic dissection of human neurodevelopment, enabling transformative insights from the study of genetic and environmental causes of neuropsychiatric disorders, a community-wide effort to which we and several others have been contributing ^2,5–14^. Importantly, our recent benchmark of cortical brain organoids (CBO) compared to the human fetal cortex confirmed the preservation in CBO of transcriptional programs pinpointed as relevant for disease modelling ^15^.

Despite these advances, the characterization of brain organoids at single cell resolution from entire cohorts, and in perspective at population-scale, remains however an unmet challenge, though an obviously required one if we are to capture how individual genomes and developmental trajectories shape variability in vulnerability and resilience across the spectrum of neurodiversity ^16–18^. Scaling up human brain organoids modelling and molecular profiling by single cell omics would allow to understand how the molecular causes of neurodevelopmental disorders (NDDs) trigger deviations from physiological trajectories ^19^, in line with the expanding set of population-level single cell studies ^20,21^. This is however still an experimental and analytical challenge due to high cost and workload, and to the inherent batch-to-batch variability of the complex experimental designs required.

To overcome some of these problems, progress has been made in single cell multiplexing strategies, including methods based on samples barcoding ^22–26^ and methods leveraging the detection of natural genetic variants ^27–30^. These approaches have indeed proven to be instrumental for population genetics and disease modelling studies, including through co-culture of cell lines derived from multiple donors in a single dish ^31–38^.

However, multiplexing has not yet been systematically applied to organoids, a challenge that is particularly relevant for the brain given the long term longitudinal unfolding of highly heterogeneous combinations of cell types, and the field still lacks studies able to establish the experimental and computational viability of applying multiplexing strategies to complex 3D experimental systems.

We thus implemented and benchmarked complementary strategies to multiplex human brain organoidogenesis *in vitro*, pooling induced pluripotent stem cells (iPSCs) coming from different individuals either during organoids generation, termed mosaic model according to standardised guidelines for brain organoids nomenclature we recently contributed to ^39^, or before single cell-RNAseq library preparation. To improve genetic-based cell identification when dealing with brain organoid single cell transcriptomes, we developed a *in silico* deconvolution method (SCanSNP), which we benchmarked against existing deconvolution tools, producing a consensus pipeline for robust genotype identification across datasets of different quality. Finally, we evaluated the two multiplexing paradigms through a deep reconstruction of neurodevelopmental trajectories and providing proof of principle of their suitability for linking genetic variation to neurodevelopmental trajectory phenotypes. This provides the community with an enabling resource for scaling up brain organoid modelling to the challenges of human neurodiversity.

## Results

### Multiplexing strategies for longitudinal single cell transcriptomics profiling of cortical brain organoids from different individuals

To test the feasibility of multiplexed brain organoid modelling over extended developmental time courses, we developed an experimental design comparing two approaches with distinctive features in terms of scaling, standardisation potential and experimental challenges: i) pooling iPSC lines from multiple donors to generate mosaic cortical brain organoids (mCBO, also referred to as upstream multiplexing); and ii) generating CBO individually and pooling them only prior to single cell droplet encapsulation (hereafter referred to as downstream multiplexing, see methods for details about specific library preparation used for each sample) (Fig. 1A). For the first strategy, we pooled iPSC lines (5000 cells/line) prior to organoids generation and longitudinally profiled the resulting mCBO through single cell-RNAseq at 50, 100 and 300 days of differentiation, following the same protocol previously used and benchmarked in our laboratory ^5,11,15^. For downstream multiplexing, CBOs were generated individually from the iPSC lines and profiled at the same time points, pooling equal amounts of cells per line after organoid dissociation for single cell library preparation. For both approaches, individual cell identity was demultiplexed through genetic variation (Fig. 1B).

**Fig. 1.**
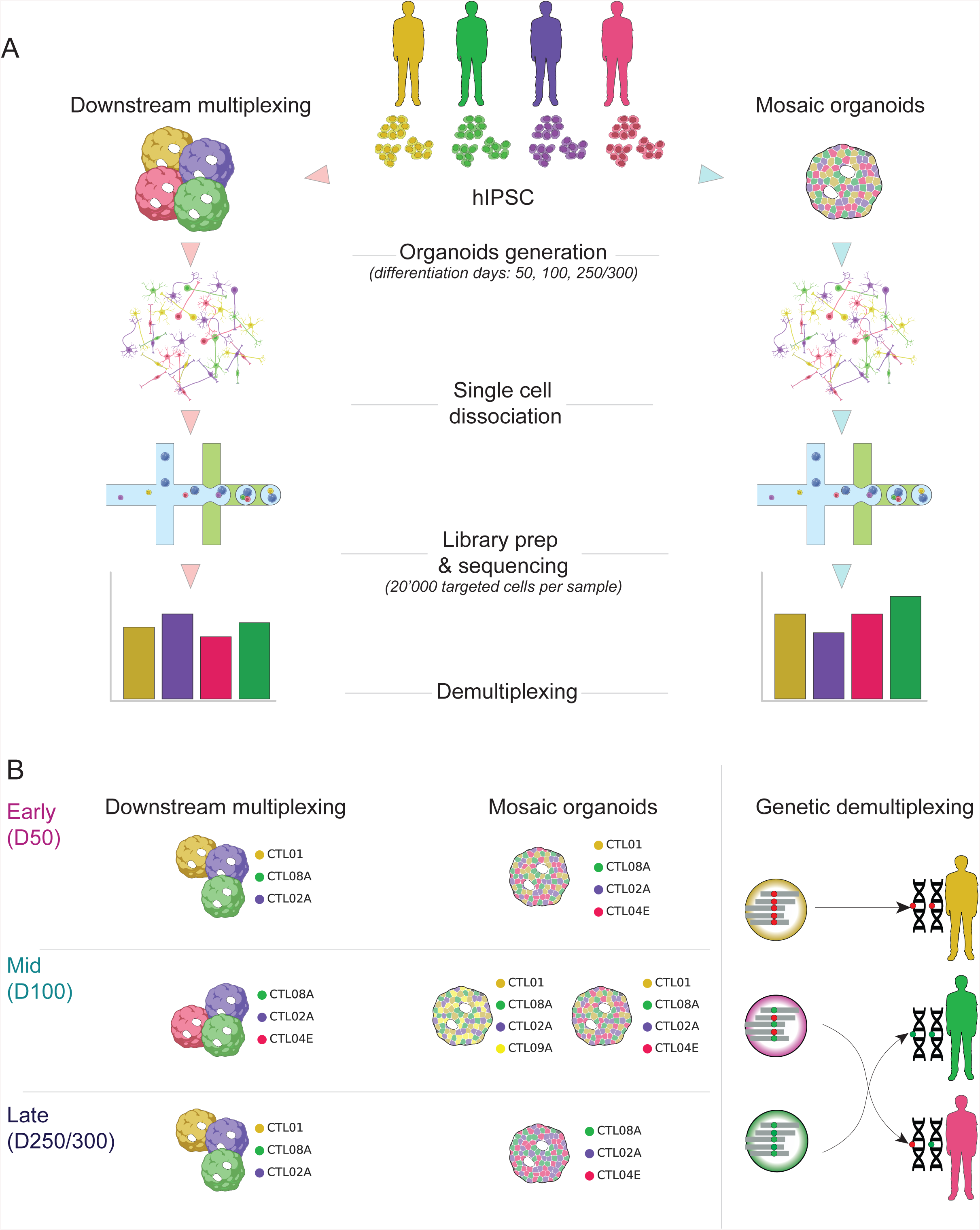
Schematic representation of Multiplexing paradigms and experimental design. **A)** Representation of the two explored multiplexing paradigms. Left side: downstream multiplexed organoids grown from individual lines and pooled in equal amounts after dissociation at the single cell level; right side: upstream multiplexed (mosaic) organoids generated by pooling equal amounts of multiple hiPSC lines during organoids seeding. After single cell dissociation both paradigms undergo library preparation, sequencing and demultiplexing. **B)** Representation of experimental design and demultiplexing approach. The panel shows the CBOs differentiation timepoints, the number of replicates for each time point and their division between the two multiplexing paradigms and the iPSC lines (genotypes) used for each experiment.

### Experimental assessment of mosaic cortical brain organoids

We first characterised mCBOs by immunofluorescence for canonical markers of neurodevelopment previously defined for organoid differentiated with the same protocol by us and others ^15,39,40^, and confirmed their expected expression patterns (Fig. 2 A,B). Next, to estimate the stability of the presence of the different iPSC lines within mCBO throughout development, we used a two-pronged approach. First, we assessed longitudinally the proliferation rate of organoids derived from individual lines in two differentiation replicates through flow cytometry-based cell cycle analysis. This showed similar proliferative trends across lines, with the expected decrease of proliferation along differentiation, and no significant variations in the proliferative rate of the different lines over time points (Fig. 2C,D Supplementary Table 1). Next, we probed the extent to which individual lines, despite homogeneous proliferation rates in isolation, could still yield skewed distributions when grown as mosaics. In line with *in vivo* evidence of high asymmetric clonality during human brain development ^41^, the quantification of genotypes from single cell transcriptomics confirmed a variable degree of balance between individual lines during mCBO differentiation (Fig. S1E).

**Fig. 2.**
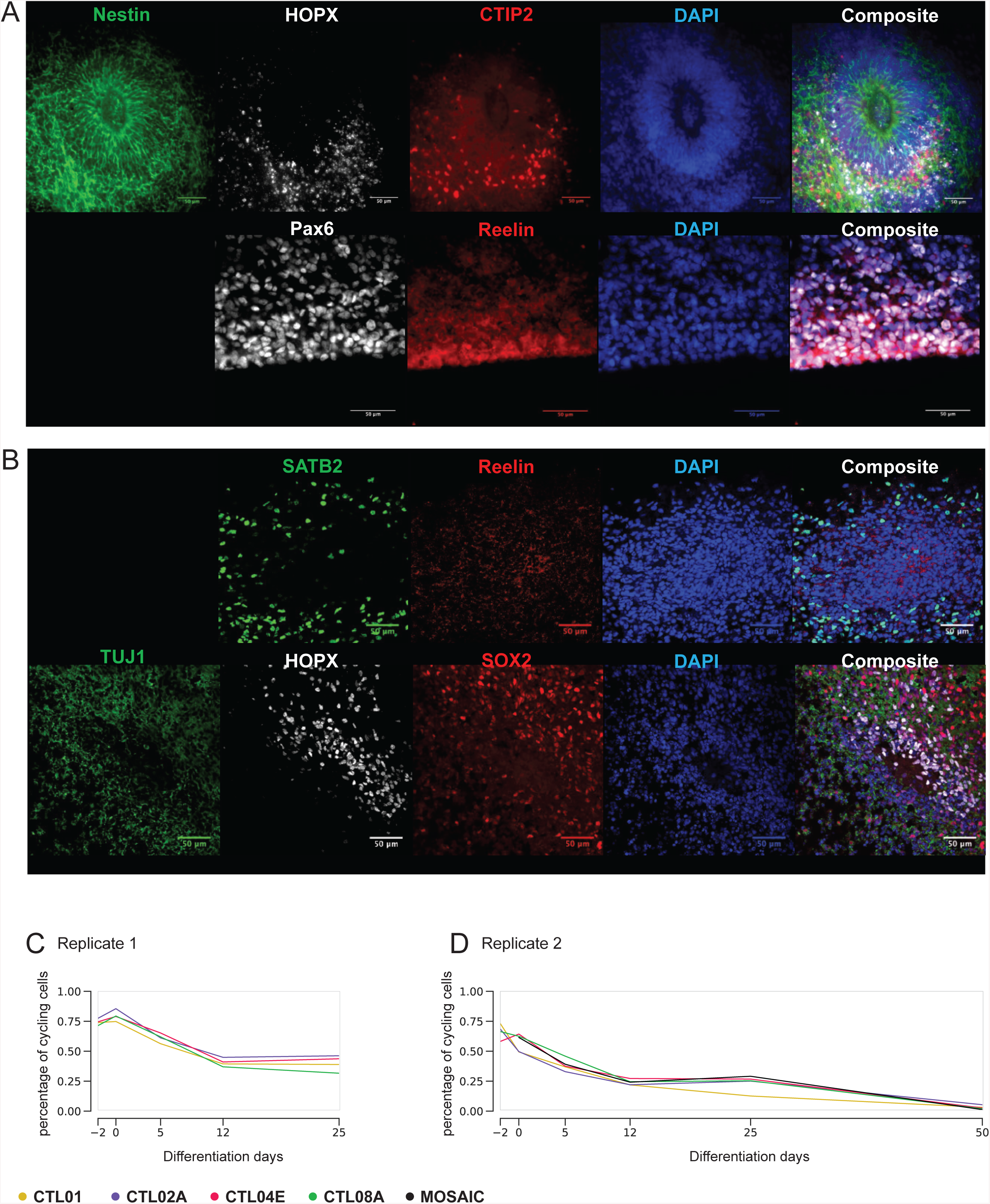
Experimental assessment of mosaic cortical brain organoids. **A). B).** Immunofluorescence-based benchmarking of neurodevelopmental markers in mosaic CBOs. At differentiation day 50 **(A)**, mosaic CBOs show consistent presence of ventricular-like structures positive for the neural stem cell marker Nestin and surrounded by a layer of cells expressing the outer radial glial marker HOPX. The presence of newborn deep-layer, CTIP2+ (BCL11B) cortical neurons can be appreciated next to HOPX+ cells. On the outer surface, mosaic CBOs reveal the presence of Reelin+ cells, consistent with the presence of Cajal-Retzius cells. At differentiation day 100 **(B)**, mosaic CBOs display more mature ventricular structures characterised by the presence of HOPX+ outer radial glial cells, SATB2+ intermediate layers neurons and SOX2+ neural precursors. Weak Reelin signal is detected and is supposedly due to the secretion of the chemotactic protein by more superficial regions of the organoids. **C)** and **D).** Percentage of cycling cells in pure lines and mosaic organoids detected by FACS (methods) performed on two differentiation replicates at relevant timepoints, from day −2 (organoids seeding) to day 25 **(C)** and 50 **(D)**.

### Benchmarking of demultiplexing algorithms

The single cell transcriptomics datasets generated from both upstream and downstream-multiplexed organoids at early, mid and late stages of differentiation (Fig. 1B) were demultiplexed using three different state of the art algorithms available for genetic demultiplexing (Demuxlet ^27^, Souporcell ^28^, Vireo ^42^). We observed an unexpectedly high number of predicted doublets, with a puzzling count distribution between singlets and doublets (Fig. S1 A,B). This was associated with a variable degree of unassigned cells and agreement among tools (Fig. S1 C,D). We thus set out to develop a new method, SCanSNP (Fig. 3A), to overcome the observed limitations by: i) dividing the classification challenge into two steps, one for identity assignment and one for doublets detection, and ii) measuring genetic purity of each droplet to identify low-quality droplets, and separate them from authentic doublets. Finally, to consolidate deconvolution accuracy, we set a consensus call framework considering strength and weaknesses of each algorithm, including non-genetic based tools for doublets and low quality cell detection ^43,44^, merging their outcomes into one combined call (Fig. 3B). Next we performed again the demultiplexing step using all the above approaches. We found that all three so far available tools tend to overestimate doublets, if compared to the theoretically expected rate, SCanSNP or the consensus call (Fig. 4A). In particular, SCanSNP and the consensus call were the only ones for which the average log-counts distribution of UMIs on predicted doublets resulted, as expected, higher than singlets (Fig. 4C). The reason is likely due to the fact that existing algorithms exhibit a bias in doublets detection that includes low quality droplets, that SCanSNP is instead able to identify (Fig. S1C). Accordingly, the evaluation of agreement rate in the assignment of individual identity across algorithms highlighted a high overall agreement for singlets’ calls, with cases of lower agreement coinciding with datasets that had higher genotype imbalance (Fig. S1D). On the other hand, we observed that doublets detection agreement was consistently lower (Fig. S1D). Thus, we also assessed the demultiplexing performance of the different algorithms using 5 *in silico* multiplexed single cell datasets to benchmark them against a ground-truth reference spanning datasets with varying degrees of balance between genotypes. As shown in Fig. 4B, all demultiplexing algorithms performed better in balanced cases and SCanSNP resulted to be the most accurate in both balanced and imbalanced settings.

**Fig. 3.**
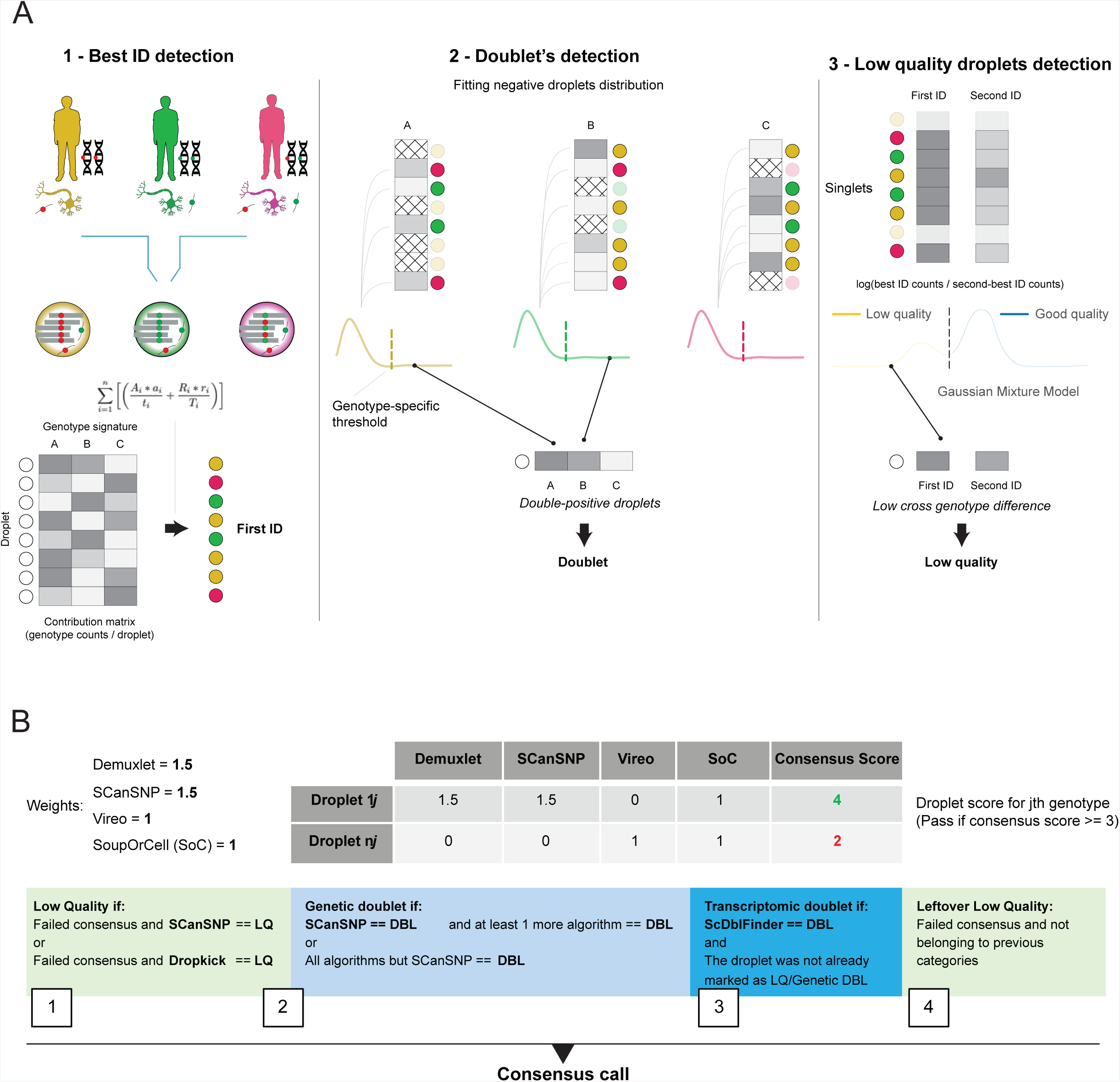
SCanSNP and consensus call overview. **A)**. Main steps of SCanSNP: 1) Best ID assignment, 2) Doublet’s classification, 3) low quality droplets detection. Colours represent different individuals and droplets identity after best ID assignment, grey-scale intensity in matrices represents the number of reads, cross-patterned cells represent droplets not included in the computations, coloured curves represent fitted distributions. **B)** Schematic representation of consensus call. The top section illustrates an example of consensus score with weights for each algorithm. Weights are the partial score attributed from different algorithms towards a specific genotype. The bottom section shows how consensus, genetic and non genetic software outputs are combined to define singlets, doublets and low quality cells. DBL and SoC are abbreviations for doublets and Souporcell respectively.

**Fig. 4.**
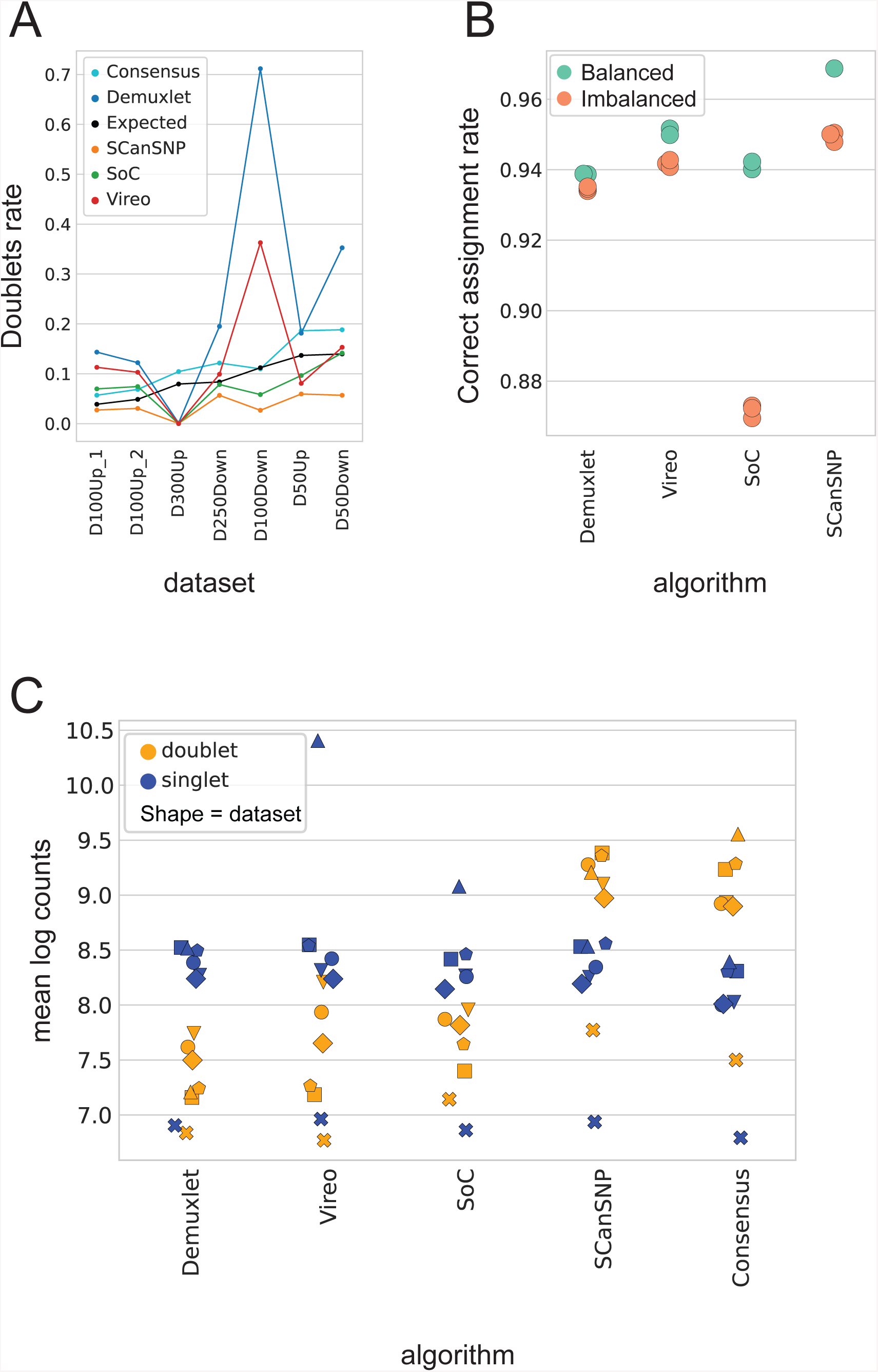
SCanSNP and consensus call benchmark. **A)** Percentage of doublets detected from each algorithm across datasets. x-axis: datasets ordered by the number of retrieved cells; y axis: doublet’s rate. Lines are coloured by algorithm. **B)** Benchmark of demultiplexing performance against ground truth in *in silico* multiplexed dataset. X axis reports different algorithms while y-axis reports the deconvolution accuracy as rate of barcodes with predicted IDs that match ground truth vs total detected barcodes per dataset. The benchmark was performed on 5 *in-silico* datasets generated from five independent single cell datasets already in house (methods). The individual experiments were multiplexed *in silico* to have balanced (light green) or imbalanced genotypes (orange). Plot is bounded between .86 and .98 to magnify the differences among and within tools, and jittered on y axis for readability. **C)** Distribution of predicted singlets and doublets from the different algorithms for each of the 7 datasets. Algorithms are shown in *x* axis, and mean log counts on y axis. Shape of each symbol corresponds to a dataset. Markers are colored by singlet or doublet predicted labels. SoC is abbreviation for Souporcell

### Single cell analysis of neurodevelopmental cell types in cortical brain organoids

After deconvolving the identity of each droplet of our dataset by the consensus call, we discarded doublets and low-quality cells, and proceeded with cells coming from the iPSC lines across multiplexing modalities (Fig. S1E, Supplementary Table 6). We thus analysed the single cell transcriptomic dataset entailing 4 iPSC lines, 3 time points and 2 multiplexing modalities (Fig. 1B). We pre-processed each sample individually, integrated them and carried out multi-tier filtering. Force-directed graph, a dimensionality reduction technique that highlights the most differentiated cell populations (the pointed extremities of the graph) while preserving local differences, was used to visualise and analyse the integrated dataset (Fig. 5A).

**Fig. 5.**
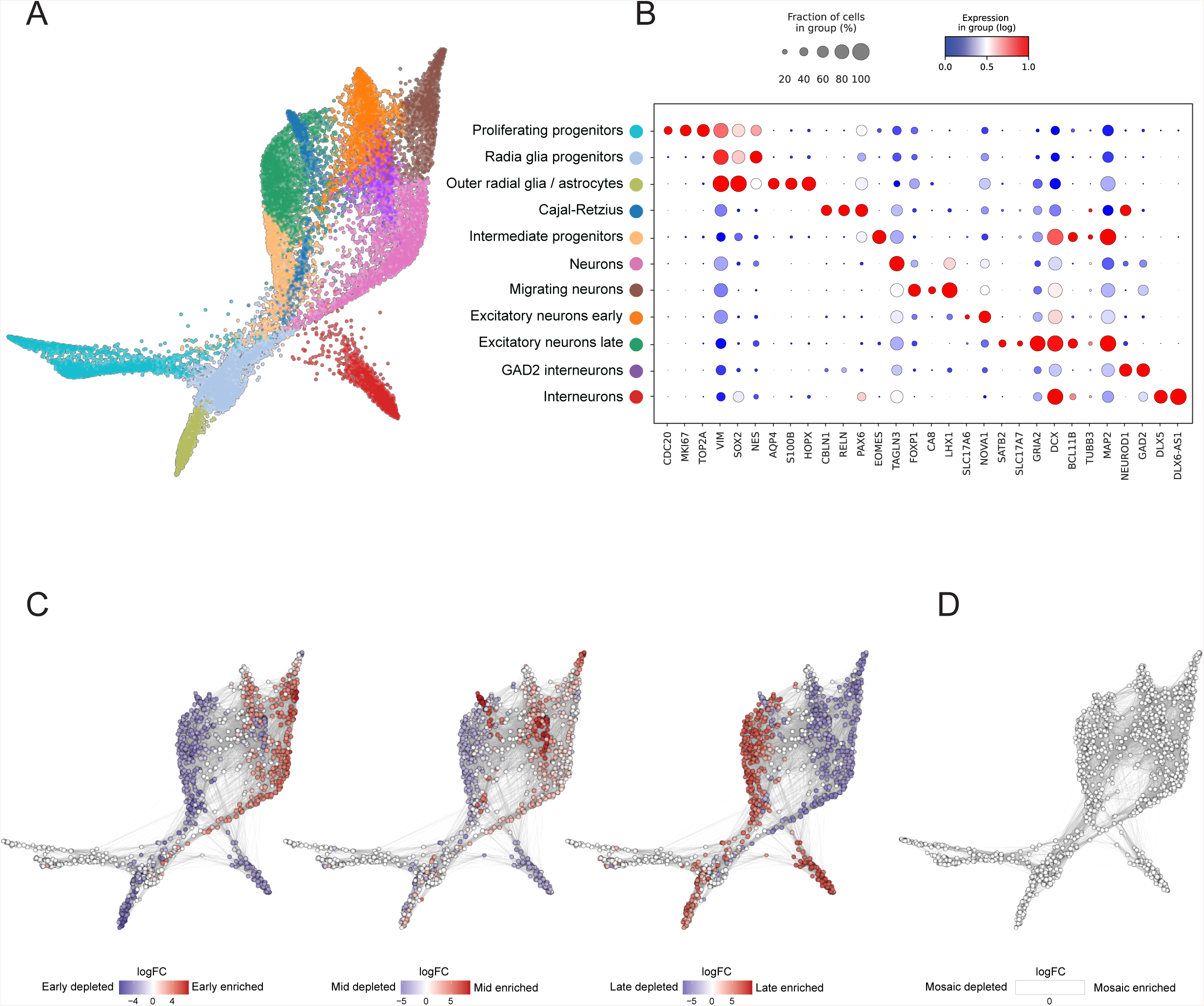
Sample multiplexing allow identification of stage specific neurodevelopmental cell populations. **A)** Cells embeddings inforce-directed graph after pre-processing and filtering. Each dot is a cell, coloured by annotated cell type. **B)** Dotplot of gene expression for some of the relevant markers used in the annotation. Size of dots is proportional to the number of cells expressing a marker and colour encodes the mean expression in group (lognorm counts). **C)** Differential abundance graph of CBOs data across timepoints. The basis for the visualisation is the force directed graph. Dot represents groups of similar cells, size encodes number of cells in each group, line size shows the number of common cells among groups, colour code indicates enrichment for indicated categories (spatial FDR < 0.1). **D)** Differential abundance graph of CBOs data across multiplexing modalities. The basis for the visualisation is the force directed graph. Dot represents groups of similar cells, size encodes number of cells in each group, line size shows the number of common cells among groups, colour code indicates enrichment for indicated categories (spatial FDR < 0.1).

We systematically annotated the identity of each cell population by analysing the expression distribution of relevant markers, defining the gene signatures driving each cluster, and projecting the cells onto a reference single cell human fetal brain dataset (Fig. 5A, S2B, S2C, Supplementary Table 2) ^45^. Consistent with our previous benchmarking ^15^, and in line with evidence from recent studies characterising brain organoids at single cell resolution ^46–48^, CBOs recapitulated most of the relevant neurodevelopmental cell populations of human corticogenesis. We observed a well-defined group of cells expressing the proliferative markers CDC20, MKI67, TOP2A, that were labelled as proliferating progenitor cells. Those cells are in continuity with a bigger cluster of cells expressing PAX6, VIM, SOX2 and NES annotated as radial glia. This cluster undergoes a bifurcation, extending either towards cells expressing outer radial glia (oRG) / astrocytes markers HOPX, AQP4, S100B, or neuronal ones STMN2, DCX, GAP43, which we annotated accordingly. We then further divided the neuronal branch into intermediate progenitors, expressing EOMES, two clusters of excitatory neurons expressing NEUROD2, TBR1, SATB2, SLC17A6, SLC17A7, SLA, GRIA2, two interneuron clusters, expressing DLX2, DLX5, DLX6-AS1, GAD1 and GAD2, one cluster of migrating neurons, expressing FOXP1, CA8 and LHX1, and one cluster expressing RELN, PAX6 and CBLN1, annotated as Cajal Retzius cells (Fig. 5 A,B, Fig. S2B).

We then compared the different timepoints of our longitudinal dataset observing, as expected, differences in cell type abundance. While radial glia progenitors, proliferating progenitors and maturing neurons were evenly distributed across timepoints, we observed that migrating neurons, early excitatory neurons, and Cajal Retzius cells were more abundant in the early and middle timepoints, whereas oRG/astrocytes, late excitatory neurons and interneurons were more abundant in the late timepoint (Fig. S2D). Instead, the abundance of cell type proportion between upstream and downstream multiplexed samples was comparable (Fig. S2D). We thus tested for statistical significance in differential abundance, leveraging MILO ^49^, which models the differences in the abundance of cell states among experimental conditions using graph neighbourhoods and thus overcomes the pitfalls of using discrete clusters. This analysis confirmed our observations highlighting significant differences only across timepoints (Fig. 5C,D) and showing no significant difference between upstream and downstream multiplexing paradigms.

Finally, to further probe the transcriptional regulation of specific cell types in the CBOs vis a vis the *in vivo* counterparts, we took advantage of our multi-genotype dataset to detect loci with allelic-specific expression (ASE, ^50^) since the likelihood of observing heterozygous loci is higher. As expected, the number of detected ASE correlates with the total amount of reads (Fig. S2E). Moreover, exploring the correlation of proportion of reads in each allele across cell types, we observed that also in CBOs the patterns of ASE are cell type specific, in line with external evidence from human brain ^51^ (Fig. S2F).

### Multiplexed Cortical Brain Organoids recapitulate key neurodevelopmental trajectories

One of the most powerful innovations enabled by single cell analysis of experimental models featuring extensive cellular heterogeneity, such as CBOs, is the possibility to analyse developmental trajectories through pseudotime analysis ^52,53^. To deeply explore this aspect in our dataset we leveraged Partition-based graph abstraction (PAGA, Fig. S3A) ^54^ and isolated all biologically relevant lineages recapitulated in CBOs (progenitors −> excitatory neurons; progenitors −> interneurons; progenitors −> astrocytes; progenitors −> Cajal-Retzius cells; progenitors −> migrating neurons). We could thus analyse through diffusion pseudotime (DPT) ^55^ how the single cells from the different timepoints and multiplexing modalities were distributed along those neurodevelopmental trajectories (Fig. 6A,B). Moreover, we harnessed the power of trajectory-based differential expression based on generalised additive models (tradeSeq ^56^) and identified the key genes driving each lineage of differentiation (Fig. S3B, Supplementary Table 3). Noteworthy, for the excitatory lineage, trajectory analysis allowed us to identify two temporally divergent paths of neurogenesis that reconcile towards the end into the excitatory neurons clusters, where EOMES was one of the key drivers of the difference (Fig. 6A). To our knowledge this is the first empirical demonstration of the ability of CBOs to recapitulate the developmental stages of the two different patterns of human cortical neurogenesis, represented by the asymmetric divisions during early stages that generate ’self-renewing’ radial glia alongside daughter cells that go on to become neurons, and a second mechanism that involves the production of intermediate progenitor cells, that *in vivo* migrate to the embryonic subventricular zone where they produce neurons through symmetric divisions ^57–59^.

**Fig. 6.**
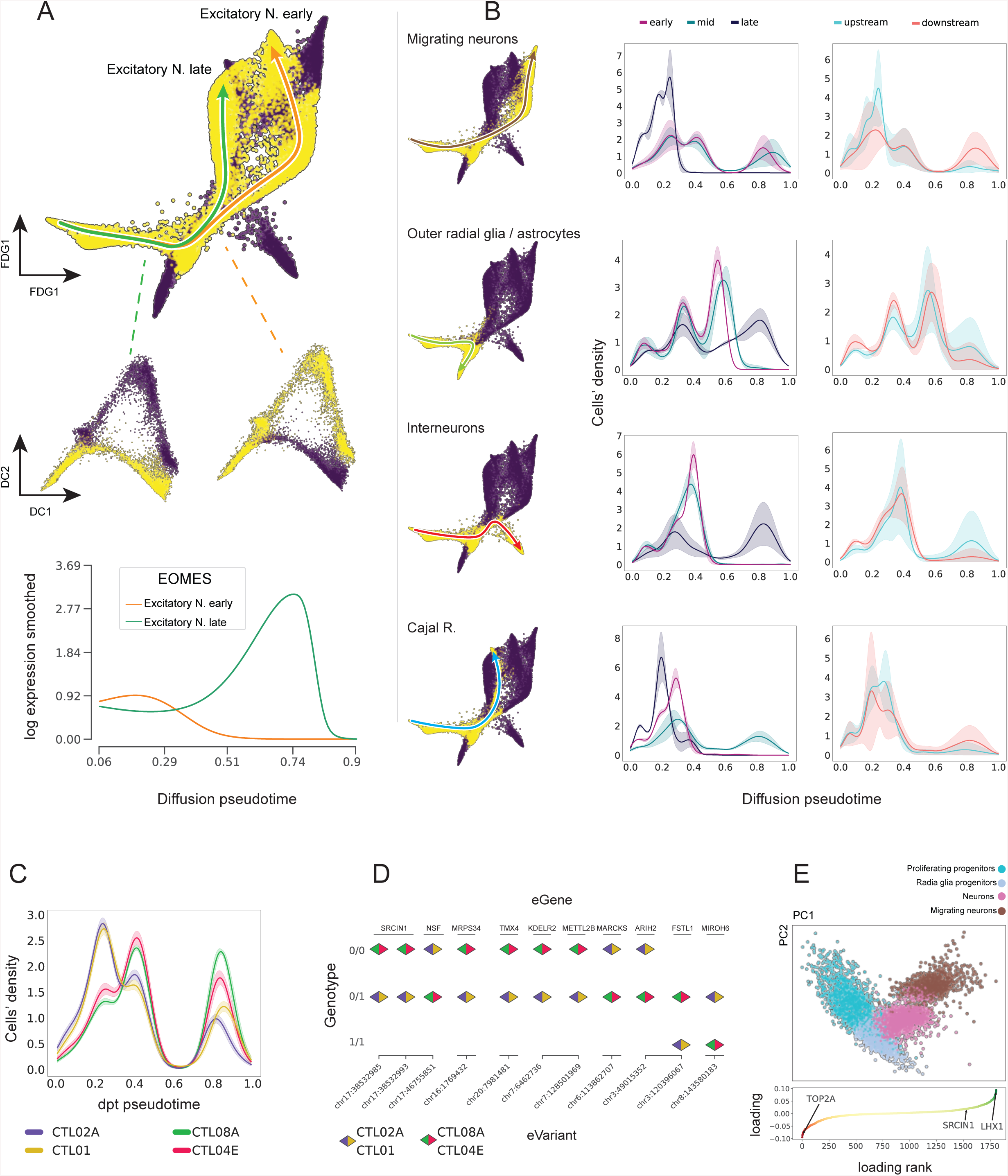
Multiplexed Cortical Brain Organoids recapitulate key neurodevelopmental trajectories. **A)** Highlight of late and early Excitatory neurons l branches. Top panel shows embeddings in force-directed graph, the middle one is a magnification of isolated branches on diffusion map, the lower panel shows the smoothed expression of EOMES for each of the 2 trajectories along pseudotime. **B)** Highlight of main developmental lineages on force-directed graph. Next to Each draw graph density plots display the number of cells along pseudotime by i) differentiation timepoints, and ii) multiplexing paradigm. Faded colour shows 1 standard deviation among dataset replicates **C)** Density plot of cells along pseudotime for the isolated migrating neurons lineage. Cells are divided and coloured by genotype. Faded colour shows 1 standard deviation across random subsampling iterations **D)** Schematics of SNPs classified as eQTLs from *jerber et al. 2021*, and their allelic configuration (Ref/Ref, Ref/Alt, Alt/Alt), for the pairs of genotypes exhibiting difference in pseudotime. **E)** PCA (harmony integrated) of isolated migrating neurons branch The lower panel displays migrating neurons-specific highly variable genes. Genes are ordered by their PC1 loading, with TOP2A (Cycling progenitors marker), LHX1 (Migrating neurons marker), and SRCIN1 (only eGene from D also present in highly variable genes) highlighted.

### Linking genetic variation to the cellular processes underpinning human neurodevelopment

Genome-wide association studies (GWASs) have uncovered hundreds of thousands of genetic variants associated with human traits ^60^. Mechanistically linking such variants to phenotypes of interest by molecular quantitative trait locus (molQTL) mapping has been instrumental to understand functional effects of genetic variation ^61^. In particular, single cell expression quantitative trait locus (eQTL) modelling allows to identify the impact of single nucleotide polymorphisms (SNPs) on cell-type-specific molecular mechanisms, as we pioneered in ^62^, pushing forward previous work by the GTEx Consortium to characterise variation in gene expression across individuals and diverse tissues of the human body ^63^. Moreover, since single cell analysis can quantify longitudinal trajectories, it is possible to assess the effects of SNPs on traits that vary dynamically along a continuous axis ^20,64^. The power of these approaches however is yet to be exploited for developmental variation as a trait per se that is now amenable to longitudinal quantitation in the form of single cell resolved trajectories. We thus set it out to leverage our *in vitro* cohort and probe as proof-of-principle how genetic variants could be linked to the neurodevelopmental traits quantified in CBOs. First, we empirically tested in our dataset the statistical power to perform single cell eQTL analysis through our previously developed tool, scPower ^65^ and identified the number of lines needed to perform single cell *de novo* eQTL discovery, given the CBO-specific variance we had determined in terms of gene expression across neurodevelopmental cell types (Fig. S3C). Next, we followed a supervised approach to investigate the genetic basis of the differences we observed in neurodevelopmental trajectories between individuals during CBOs differentiation.

In particular, we focused on the trajectory of migrating neurons since it displayed the greater genotypes’ diversity in the distribution of cells along pseudotime (Fig. 6C, S3D). Upon downsampling cells per genotype to ensure balance within each timepoint, we found differences between 2 subgroups (of 2 individuals each) (Fig. 6C, Supplementary Table 4). We identified the genetic variants that distinguish the 2 subgroups and found that 11 of these genetic variants (Fig. 6D, Supplementary Table 5) have been previously identified as single cell eQTL in a population-scale single-cell RNA-seq profiling of iPSC derived neurons (see *Jerber et al*. 2021 ^35^ Table 7), with two of them marked as cis-acting eQTLs for SRCIN1, one of the trajectory-specific highly variable genes we had identified (Fig. 6E), that encodes for a protein previously shown to be a regulator of disease relevant alterations in cell migration ^66,67^ and hence potentially representing the genetic underpinning of the neurodevelopmental difference in the migrating neurons trajectory observed here between the CBOs from different individuals.

## Discussion

In this work we tackle the multiplexing need to leverage brain organoids modelling for the systematic study of human neurodiversity. Pooling strategies already proved transforming for iPSC-based disease modelling ^34–38^, however their application to the inherently complex setting of brain organoids, which generate a highly heterogeneous composition of cell types over very extended times, had not yet been undertaken.

We showed that different iPSC lines follow similar trends of cell proliferation rates along CBOs differentiation and can be multiplexed in mosaic CBOs that recapitulate the expected cytoarchitectural organisation, including ventricular-like structures and the gradual emergence of relevant neurodevelopmental markers of standard CBOs, with an expectedly variable degree of balance between the individual lines was observed.

The guidelines emerging from this work thus indicate to opt for the downstream multiplexing strategy when the biological question at hand requires strict balance among individuals, with the mosaic model that is instead more suited for unbiased large scale studies, in the same way as developmental neuroscientists can leverage pooled CRISPR screening systems for screening the impact of genetic variants, in combination with validations of single mutations ^68–71^.

Furthermore, it will be useful for the field to further characterise the spatial distribution of lines in mosaic CBOs, as well as to study the developmental features of mosaic CBOs using higher number of iPSC lines and/or iPSC lines reprogrammed from patients and neurodiverse individuals.

The unique opportunity to deconvolute individual identity from single cell transcriptomics data leveraging genetic variability, allowed us to use mosaic CBOs to study molecular processes of human neurodevelopment at high-resolution and at increased scale. Here we benchmarked and improved the demultiplexing steps leveraging and aggregating the pros of different publicly available state of the art algorithms. Moreover we developed SCanSNP, a new deconvolution method that more accurately identifies doublets and low quality cells, and is flexible in case of imbalanced genotypes.

We next deeply characterised the neurodevelopmental cell types and trajectories recapitulated by CBOs across timepoints and multiplexing modalities. Noteworthy, the unicity of our longitudinal design allowed us to capture a very interesting and relevant neurodevelopmental process represented by the transient emergence of Cajal Retzius cells, a population that was also recapitulated in a different brain organoids model ^47^. In our dataset, Cajal Retzius neurons are enriched at middle stages and then depleted at advanced stages, in both downstream and mosaic CBOs (Fig. 4 and Fig. S4) consistently with *in vivo* evidence that they are fated to elimination as neuronal networks mature ^72^.

This design also allowed us to identify, for the first time at the level of single cell analysis of cortical brain organoids, the developmental divergence of direct and indirect neurogenesis, highlighting EOMES as the key driver guiding the different paths of early versus late glutamatergic neurons differentiation in the organoids, thus confirming the ability of this experimental model to recapitulate the physiological processes of a unique aspect of human neocortical development, represented by the spatiotemporal regulation of neural intermediate progenitor cells that serve as a basis for interpreting human brain development, evolution, and disease ^57^. At the current resolution afforded by the single cell transcriptomic interrogation of CBOs, this domain of indirect neurogenesis is likely capturing the heterogeneous populations of iNPC recently described from in vivo datasets, whose further partition remains an important challenge for further improving organoids capture of in vivo trajectories ^57,58^.

Moreover, the results from human samples suggest that EOMES might be capable of promoting inhibitory neuronal identity as well ^57^. In our cortical brain organoids, even if patterned for the dorsal forebrain ^73^, we observed the differentiation of GABAergic neurons at late stages, confirming the evidence that human cortical progenitors are also capable of producing GABAergic neurons with the transcriptional characteristics and morphologies of cortical interneurons ^74^ (in contrast to previous developmental studies in rodents that had led to a prevailing model in which cortical interneurons are born from a separate population of progenitors located outside the developing cortex in the ganglionic eminences) and the suggestion that inhibitory neurons in humans may arise from intermediate progenitors in the cortex at late stages of neurogenesis ^57^.

Through the analysis of developmental trajectories, we showed how brain organoid multiplexing can be used to link genetic variants to specific and dynamic neurodevelopmental phenotypes, captured in particular along the transcriptional regulation of migrating neurons. This paves the way for molQTL discovery once multiplexing approaches will be applied to CBOs differentiated from large scale cohorts of iPSC lines.

Indeed we showed that the number of lines needed for *de novo* eQTL discovery (which we propose to term eDTL from expressed Developmental Trait Loci) should be in the range of hundreds of individuals (Fig. S3C), which indicates that an automation-based leap in throughput is needed before that longitudinal organoid series from actual population-scale cohorts can be mined for that.

Finally, the mosaic model represents a transforming tool to study gene–environment (GxE) interactions by allowing environmental chemicals ^11^ or drugs to be studied on multiple genetic backgrounds ^75^, thereby pushing forward precision neurotoxicology and pharmacology.

## Methods

### Culture of hiPSCs

hiPSC lines were cultured under feeder-free conditions on matrigel coated plates at 37 °C, 5 % CO2 and 3 % O2. To coat culture dishes, matrigel solution was prepared by diluting matrigel (Corning, #354277) 1:40 in ice-cold DMEM/F12 (Gibco, #11330057) and stored at 4 °C until use. Before plating cells, 6cm dishes were coated with 1 mL of matrigel solution and incubated for 30 minutes at 37 °C. hiPSCs were maintained in TeSR/E8 medium (Stemcell Technologies, #05990) supplemented with 100 U/mL penicillin and 100 ug/mL streptomycin (ThermoFisher, #15140122) with daily media changes and passaged 1:8 to 1:10 when confluency reached around 70%. To detach cells, plates were rinsed with 2-3 ml of PBS (Gibco, #10010023) and treated with 0,5 mL ReLeSR (Stemcell technologies, #05872) for 5 minutes at 37 °C. When single cell dissociation was needed, Accutase (Sigma-Aldrich, #A6964) was used instead of ReLeSR, and 5uM ROCK inhibitor Y-27632 (Tocris, #1254) was added to the medium to enhance cell survival in the first 24 hours. All subjects signed an informed consent and the use of hiPSCs was approved by the ethical committee of the University of Milan. All hiPSC lines were reprogrammed by at least 15 passages and verified to be mycoplasma free by routine PCR testing.

### Multiplexing strategies

In the experimental design we adopted two distinct multiplexing strategies, namely upstream and downstream, that differ for the moment in which the different cell lines are mixed.

In the upstream multiplexing approach, cortical brain organoids were generated by mixing equal amounts of PSCs derived from each cell line to obtain mosaic brain organoids. Briefly, hiPSC lines were dissociated in parallel at the single cell level, cells were counted separately and then mixed in equal proportions, (namely 25% from each line) to obtain a mosaic cellular suspension. After diluting the cell suspension to the desired concentration of 2×10^5 cells/ml, organoids were generated as explained in the dedicated chapter.

In the downstream approach, cortical brain organoids were independently generated from each individual hiPSC line and grown separately. When reaching the desired analytical timepoint, organoids from each cell line were dissociated in parallel, cells were counted and mixed in equal amounts to obtain a pooled cell suspension.

For the comparison of neurodevelopmental cell types and trajectories between upstream and downstream multiplexed CBOs the design included 4 hiPSC lines (CTL08A, CTL01, CTL02A, CTL04E) across replicates in both multiplexing modalities. Details about the hiPSC lines can be found in the Supplementary Table 6.

### Cortical Brain Organoids

Pure lines-derived and mosaic brain organoids were generated using an adaptation of the previously described protocol ^76^, which allows to obtain dorsal telencephalon cortical organoids, introducing orbital shaking on day 12 of differentiation as previously published by us in ^5,11,15^. hiPSCs were grown on matrigel-coated plates to approximately 60% to 70% confluency, dissociated with accutase (Sigma, #A6964) and resuspended in TeSR/E8 medium supplemented with 5uM ROCK inhibitor Y-27632 (Tocris, #1254) to reach a final concentration of 2 × 10^5 cells/ml. 100uL/well of cell suspension were seeded in ultra-low attachment, U-bottom 96 well plates (SystemBio, #MS9096UZ) and then centrifuged for 3 minutes at 150 RCF to promote formation of embryoid bodies. Plates were incubated at 37 °C, 5% CO2, 3% O2 for 2 days and then first media change was performed, substituting TeSR/E8 with neural induction medium containing 80% DMEM/F12 medium (Gibco, #11330057), 20% Knockout serum (Gibco, #10828028), Non-essential amino acids 1:100 (Sigma, #M7145), 0.1 mM cell culture grade 2-mercaptoethanol solution (Gibco, #31350010), GlutaMax 1:100 (Gibco, #35050061), penicillin 100 U/mL and streptomycin 100 μg/mL (ThermoFisher, #15140122), 7 uM Dorsomorphin (Sigma, #P5499) and 10 uM TGFβ inhibitor SB431542 (MedChem express, #HY-10431). Since that moment, defined as day 1, cultures were grown in normal oxygen conditions (5% O2). Media changes were performed daily for the subsequent 4 days and, on the fifth, neural induction medium was substituted with complete neurobasal medium, composed of neurobasal medium (Gibco, #12348017), B-27 supplement w/o vitamin A 1:50 (Gibco, #12587001), GlutaMax 1:100 (Gibco, #35050061), penicillin 100 U/mL and streptomycin 100 μg/mL (ThermoFisher, #15140122) and 0,1 mM cell culture-grade 2-mercaptoethanol solution (Gibco, #31350010) supplemented with 20 ng/mL FGF2 (Peprotech, #100-18B) and 20 ng/mL EGF (Peprotech, #AF-100-15). On day 12, organoids were transferred by pipetting with cut-end pipette tips from 96-wells to 9 cm ultra-low attachment dishes (Systembio, #MS-90900Z) and placed on a standard orbital shaker (VWR Standard Orbital Shaker, Model 1000). From day 12 onwards media change was performed every other day. On day 23, FGF and EGF were replaced with 20 ng/mL BDNF (Peprotech, #450-02) and 20 ng/mL neurotrophin-3 (Peprotech, #450-03) to promote differentiation of neural progenitors. From day 42 onwards, complete neurobasal medium without BDNF and NT3 was used, performing media changes every other day.

### Cell cycle analysis

Pure lines-derived organoids were subjected to cell cycle analysis at multiple timepoints after their generation along with cell suspensions employed in their generation. Organoids were collected at differentiation day 0, 5, 12, 25, 50 and 75, dissociated at 37 °C for 5’ or 30’ with Trypsin-EDTA (Euroclone, #ECB3042) or Papain (Stemcell Technologies, #07466) respectively according to the analytical timepoint, being papain more efficient for the dissociation of mature organoids.

As for the first analytical replicate(Fig. 2C), cells were fixed with cold EtOH and stored at +4 C. Upon completing the longitudinal cohort, cell suspensions were rinsed with PBS, stained overnight with 3uM propidium iodide solution (Thermofisher, #P1304MP) in the presence of 25ug/ml RNAse I (Thermofischer, #EN0601) and analysed with FACSCelesta (BD biosciences) to measure DNA content. Analyses were performed with FlowJo v10 software (BD biosciences).

In the second analytical replicate, to avoid the deterioration of early timepoints samples, 1 million cells per sample were resuspended in PBS supplemented with 0.1% BSA (Sigma, A9418), stained for 20’ at 37 °C with 2ug/ml Hoecst33342 (Sigma, #B2261) in the presence of 5uM Verapamil (Sigma, #V4629) and analysed with Cytoflex (Beckman Coulter life sciences) to measure DNA content.

Analyses were performed with FlowJo (Replicate 1, BD Biosciences) or FCS-Express 7 software (Replicate 2, DeNovo software). Raw data are shown in Supplementary Table 1.

### Histological analysis

Organoids were collected on differentiation days 50 and 100 washed in PBS and fixed overnight at 4 °C in 4% paraformaldehyde/PBS solution (SantaCruz, #sc-261692). After rinsing with PBS twice, samples were embedded in 2% low melting agarose, put in 70% v/v ethanol and immediately given to the facility for paraffin encasing, sectioning and routine haematoxylin/eosin staining.

Deparaffinization and rehydration was achieved by consecutive passages of 5 minutes each in the following solutions: 2x histolemon (Carlo Erba, #454912), 100% ethanol, 95% ethanol, 80% ethanol and 2x ddH2O. Sections were then incubated for 45 min at 95 °C with 10mM Sodium citrate buffer (WVR chemicals, #27833.294P) + 0,05% Tween 20 (Sigma, #P1379) for simultaneous antigen retrieval and permeabilization; then equilibrated at RT for at least 2 hours. After 30 minutes blocking with 5% normal donkey serum (Jackson ImmunoResearch, #017-000-121) in PBS, incubation with primary antibodies in PBS + 5% normal donkey serum was performed. The following primary antibodies were used: Pax6 (Rabbit, 1:200, Biolegend), Tuj1 (Mouse, 1:1000, Biolegend), Nestin (Mouse, 1:500, Millipore), Reelin (Mouse, 1:400, Millipore), CTIP2 (Rat, 1:400, Abcam), SATB2 (Mouse, 1:400, Abcam), HOPX (Rabbit, 1:500, Sigma), SOX2 (Goat, 1:1000, R&D System). The day after, secondary antibodies conjugated with AlexaFluor 488, 594 or 647 were diluted 1:300 in PBS and applied to the sections for 1 hour, followed by 5 minutes incubation with 1ug/ml DAPI solution. After each incubation 3 × 5 minutes washing steps with TBS buffer were performed. After a final rinse in deionized water, slides were dried and mounted using Mowiol mounting solution. Images at 20X, 40X and 63X magnification were acquired using a DM6 B MultiFluo (Leica) equipped with Andor Zyla VSC-04470 sCMOS camera and then processed with FIJI software.

### Cortical Brain Organoids processing for single cell transcriptomic analysis

Organoids were collected at day 50, 100, 250 and 300 differentiation days (+/- 3 days). 3-5 organoids per condition were dissociated by incubation with a solution of 0,5 mg/ml Collagenase/Dispase (Sigma) + 0,22 mg/ml EDTA (Euroclone) with 10 ul of DNaseI 1000 U/ml (Zymo Research) for 30 – 45 min according to organoid size. Digested suspensions were filtered through 0.4 uM FlowmiTM cell strainers, resuspended in PBS and counted using a TC20 Automated Cell Counter (Biorad). For the sample D100 Down, the single cell suspension was further centrifuged at 500g for 3 min and resuspend in 200 ul of cold PBS-0,04% BSA. 800 ul of pre-chilled 100% Methanol was added dropwise, for a final 80% concentration. Cells were fixed for 30 min and stored at −80 degrees for 6 months. For recovery of single cell suspension, fixed cells were thawed at 4°C (all steps at 4°C), centrifuged at 1000g for 5 min, Supernatant was completely removed and the pre-chilled SSC cocktail (3X SSC / 0.04% BSA / 1% SUPERase-In / 40mM DTT) added. Cells were counted and resuspended to a concentration 1000 cells/ul. Droplet-based single-cell partitioning and single-cell RNA-Seq libraries were generated using the Chromium Single-Cell 3′ Reagent v2 Kit (10× Genomics, Pleasanton, CA) following manufacturer’s instructions (Zheng et al., 2017). Briefly, a small volume (6 – 8 μl) of single-cell suspension at a density of 1000 cells/μl was mixed with RT-PCR master mix and immediately loaded together with Single-Cell 3′ gel beads and partitioning oil into a single-cell 3′ Chip. The gel beads were coated with unique primers bearing 10× cell barcodes, unique molecular identifiers (UMI) and poly(dT) sequences. The chip was then loaded onto a Chromium instrument (10× Genomics) for single-cell GEM generation and barcoding. RNA transcripts from single cells were reverse-transcribed within droplets to generate barcoded full-length cDNA. After emulsion disruption, cDNA molecules from each sample were pooled and pre-amplified. Finally, amplified cDNAs were fragmented, and adapter and sample indices were incorporated into finished libraries which were compatible with Illumina sequencing. The final libraries were quantified by real-time quantitative PCR and calibrated with an in-house control sequencing library. The size profiles of the pre-amplified cDNA and sequencing libraries were examined by Agilent Bioanalyzer 2100 using a High Sensitivity DNA chip (Agilent). Two indexed libraries were equimolarly pooled and sequenced on Illumina NOVAseq 6000 platform using the v2 Kit (Illumina, San Diego, CA) with a customised paired end, dual indexing (26/8/0/98-bp) format according to the recommendation by 10× Genomics. Using proper cluster density, a coverage of around 250 M reads per sample (2000–5000 cells) was obtained corresponding to at least 50,000 reads/cell.

### Generation of bulk RNAseq-based variant calling file (VCF)

Reference genotypes for the deconvolution were obtained from bulk RNA-seq data of pure lines derived organoids.

Total RNA was isolated from pure lines organoids with the RNneasy Micro Kit (Qiagen) according to the manufacturer’s instructions. RNA quantification and integrity was assessed through Agilent 2100 Bioanalyzer electrophoretic analysis. TruSeq Stranded Total RNA LT Sample Prep Kit, Illumina was used to run the library for each sample using 500 ng of total RNA as starting material. Sequencing was performed with the Illumina NOVAseq 6000 platform, sequencing on average 35 million 50bp paired-end reads per sample.

First, raw reads were aligned on GRCh38 v93 Ensembl reference genome using STAR ^77^ two-pass mode to learn splicing junction from data, subsequently read group (RG) tags where made uniform and optical and PCR duplicated reads were marked using GATK MarkDuplicates. The resulting BAM file was recalibrated using BaseRecalibrator and ApplyBQSR with sorted 00-All.vcf.gz (b151_GRCh38p7) file from dbSNP. Next, we performed the variant calling using GATK HaplotypeCaller ^78^ with the option ‘dont-use-soft-clipped-bases’ meant to optimise the call from spliced RNA data, --standard-min-confidence-threshold-for-calling set to 30 and --min-base-quality-score equal to 20. We relied on HaplotypeCaller ‘GVCF’ mode for the call, to preserve also information of sites with homozygous reference alleles as long as the quality was acceptable, in order to retrieve the maximum cohort-wise informative power. After different trials iterations we decided to apply separated thresholds for MIN_DP, DPand quality (GQ) of 10, 10 and 20 respectively. Finally we used CombineGVCFs and GenotypeGVCFs exclusively to merge together GVCF of genotypes mixed in the corresponding SC experiment, ignoring the aggregated INFO field.

### Alignment and genotype demultiplexing

SC RNA-seq data were aligned using cellranger3.0.0 count and the matched reference provided from 10x. For subsequent demultiplexing and downstream analyses only droplets passing the cellranger filter were considered. For the demultiplexing we applied Demuxlet, Souporcell, Vireo, and SCanSNP. The final identities used are the result of Aggregated Call. With the exception of Souporcell, all the tools were provided with bulk RNA-seq derived VCF and were embedded in a collective singularity image. To optimise the deconvolution process to our samples we used Demuxlet with default parameters and setting ‘--doublet-prior’ according to the number of retained droplets after CellRanger filtering and V3 kit specs doublets expectation. CellSNP, the vireo companion tool for SC-side variant calling was launched ‘with -p 10 --minMAF 0.1 --minCOUNT 20’ and subsequently Vireo was launched on combined vcf file cleaned from loci containing missing calls. Souporcell was launched using the available singularity image specifying only the number of genotypes in the mixture (*k*). hiPSC lines included in this analysis are listed in the Supplementary Table 6.

### Single cell data preparation

All downstream analyses were performed within scanpy single cell analysis framework ^79^.

Basic filtering was done right after importing count matrices from cellranger. We inspected the number of genes, mitochondrial gene counts and ribosomal gene counts distributions and adopted dataset-specific thresholds to remove droplets with likely technical issues. Next we removed droplets called low-quality or doublets according to Aggregated call. After merging the 7 datasets, cell counts were normalised, log transformed using *sc.pp.normalize_total* and *sc.pp.log1p* scanpy functions. Finally we regressed out the effect of total counts and percentage of mitochondrial transcripts via *sc.pp.regress_out* and *sc.pp.scale* functions. All functions were run with default parameters.

### Cells filtering and Annotation

We relied on multiple tiers of annotation, filtering and dimensionality reduction. First we removed proteoglycan (WLS, ANXA2, TPBG, RSPO3, DCN, BGN, MYH3) expressing cells, considered non relevant for our downstream analyses. We next iteratively partitioned cells and assessed top marker via leiden and rank_gene_groups respectively to remove cells highly expressing adhesion (WLS, TPBG) and stress (BNIP3, PGK1, MT-CO2) markers, while manually annotating cell types using literature markers. Finally we used scanpy’s score_genes function providing er stress, and hipoxia signatures to remove clusters of cells scoring > 0 and > .3 respectively. Finally, we re-partitioned the remaining cells with leiden resolution .6 and manually transferred the annotation on new clusters, merging, if needed, different partitions into coherent cell types when key markers were overlapping.

### Highly variable genes (HVGs) selection

For HGVs detection, we took advantage of having at least 2 datasets per timepoint and detected HVGs by timepoint via *sc.pp.highly_variable_genes* scanpy function, providing each dataset as a separate batch and min_mean=0.0125, max_mean=5, min_disp=0.5 as parameters. For each timepoint we kept only genes found as HVGs in at least 2 datasets. After major filtering, and single trajectories isolation, HVGs were recomputed on the cell subset in the same fashion. Moreover we included some relevant neurodevelopment genes from literature regardless if they were detected as HVGs (Supplementary Table 2).

### Dimensionality reduction and datasets’ integration

After final dataset cleaning we computed PCA in HVGs, the subsequent datasets’ integration was carried out via scanpy’s harmony implementation ^80^ with maximum 20 iterations, cells’ neighbourhood graph was then computed on top 15 principal components (pcs), and specifying 100 as neighbourhood size (n). Upon single trajectories isolation, PCA was recomputed on each cell subset and branch-specific HVGs, harmony was run with 20 max iterations, lambda = 2, and theta = 1, cells’ neighbourhood graphs were computed with 10 pcs and 50 n, in the case of Excitatory neurons lineage we used 9 pcs and 60 n given the greater differences between early and late branch compared to within lineage continuous cells’ states of other cases.

### Trajectories isolation

To analyse the cell’s with trajectory-wise magnification, we isolated the most relevant neurodevelopmental trajectories. For this purpose we operated an higher level of cells partitioning using Leiden algorithm with doubled resolution (1.2) with respect to the original one, and used it as backbone for PAGA ^54^, obtaining a partition-based graph abstraction. The PAGA graph was refined by removing edges with weight < 0.05. After complementing the edges to 1 to obtain the equivalent distance graph we computed the shortest path from *r* root to each endpoint *e* using networkx ^81^ with Bellman-Ford method. Where *r* is the partition of cells with highest counts of “TOP2A” gene, and *e* are partitions with highest rate of the cell types considered one of the endpoints (Astrocytes, Cajal Neurons, Exc neurons early, Exc neurons late, Interneurons and Migrating Neurons).

### Differential abundance analysis

We adopted Milo ^49^ for differential abundance analysis. We tested for differential abundance between mosaic organoids dataset and non mosaic datasets via direct comparison. Subsequently tested for differential abundance between each timepoint and the other two simultaneously, therefore we provided as model.contrast *CT_n_ = T_n_ - (T_x_+T_y_)/2* for each of the *n* assessed time points, where *x* and *y* are the other two timepoints. Plots were generated displaying enriched / depleted cells’ neighbours for each timepoint with spatial FDR < 0.1.

### Developmental trajectories analysis

We aimed at assessing the distribution of cells along pseudotime by different covariates for Migrating Neurons, Astrocytes, Cajal neurons and Interneurons trajectories after their isolation. We wanted to assess whether i) timepoints differences mirror the asynchronous development of specific cell types in our *in vitro* system, ii) Multiplexing paradigm impact on the developmental timing of such populations and iii) If we had the resolution to spot differences among control genotypes. For each trajectory individually, we computed diffusion map ^82^ and diffusion pseudotime (dpt) ^55^. Next, for timepoints and multiplexing paradigms covariates we computed kernel density on each dataset (using sklearn.neighbors.KernelDensity, with) and plotted mean and +- 1 standard deviation among datasets’ densities. For genotypes’ comparison, we kept most relevant timepoints for each trajectory (early+mid for migrating neurons and Cajal neurons, early + mid + late for the other trajectories), and genotypes for which at least 50 cells were retrieved at each retained timepoint. Finally the genotypes were balanced via random sampling to have the same amount of cells across timepoints. Mean and standard deviations were computed on kernel densities of 50 sampling iterations to assess the stability.

Subsequently, we used tradeSeq ^56^ (fitGAM function was run with nknots = 8 after trimming cells in 1^st^ and 99^st^ dpt percentiles to increase stability at lowly sampled extremes) and detected pseudotime-driven transcriptional difference within each lineage for all but excitatory neurons trajectories, whereas for Excitatory neurons we used tradeSeq to find key transcriptional differences between early/mid and late neuronal trajectories.

Within lineage differences: to extract key driver genes along pseudotime we isolated highly variable genes between lineage extremes (tradeseq startVsEndTest(), pVal <= 0.001 and logFC >= 2).

Early vs Late excitatory neurons differences: to detect key differences between the two lineages we considered both the greatest divergently expressed genes at the terminal states (tradeseq diffEndTest(), pVal <= 0.001 and logFC >= 2), and the transiently varying genes, defined as simultaneously low ranked from tradeSeq:diffEndTest and high ranked from tradeSeq:patternTest(pVal <= 0.001) functions.

### Allele specific expression analysis

We leveraged our multi genotype design to carry out allele specific expression (ASE) analysis for each cell type annotated. We first produced the reads pileup at genomic (biallelic-only) loci displaying variability within our cohort using SCanSNP, which provided an anndata ^79^ with *nCells × nLoci* dimensionality and two layers: “Refreads” and “Altreads”, with the count of reads presenting or not the variant respectively. To perform the analysis, we summed-up reads mapping to variant sites (at least one heterozygous genotype) of the same cell type. If multiple genotypes were heterozygous at the given location, reads were included in the sum regardless of the original genotype. Since it is not granted that among different individuals - if present - the dominant allele is the same, we first checked that for loci with detected ASE the dominant allele was coherent prior to merging coverages from different individuals. We observed minimal discrepancy, i.e., at a given locus, the dominant allele was the vastly the same (always Ref or Alt) across genotypes. (Notebook 04_ASE/13.1_SanityCheck). For each cluster, we kept only loci with at least 20 reads, computed bValue(Alt reads / (Alt reads + Ref reads)), and performed binomial test and fdr correction (*q* < 0.05) provided by scipy ^83^ and statsmodel implementations respectively. To calculate the correlation among cell types, we used bValues of loci with detected ASE in at least one celltype and covered with at least 20 reads in all cell types. Correlation was computed in pandas with “spearman” metrics.

### SCanSNP

In our benchmark, the presence of low quality droplets and doublets were observed to be an open challenge also for well-established methods when assigning genotypes (IDs) to droplets in the genetic demultiplexing. With those challenges in mind we developed SCanSNP (available at https://github.com/GiuseppeTestaLab/SCanSNP), by breaking down demultiplexing and filtering in 3(+1) steps:

1. Best ID detection per droplet: here, as for other approaches, we leveraged the accessibility of bulk RNA-seq data to generate a function that maximises the score difference of each ID to the sequenced droplets (equation 1).

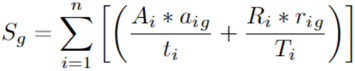

where *Sg* is the score for ID *g* in a given droplet, i are the loci for which allelic information is accessible from bulk RNA-seq, *A* and *R* are respectively the number of reads supporting Alternative and Reference alleles, *a* and *r* are the number of alternative and reference alleles in *g*, *t* and *T* are total alternative and reference alleles in the cohort at locus *i*
2. Second best ID determination: we started by creating an *m by g* contribution matrix, where *m* are the droplets and *g* are the multiplexed IDs, containing the number of reads supporting private alleles. These are alleles for which, at a given locus it is possible to trace a non-ambiguous allele-to-donor link. We used this matrix to iteratively train a multinomial logistic regression model (implemented via sklearn package), to predict which is the most likely ID after the first one, assuming ambient contamination consistent across droplets. We split the contribution matrix into groups of droplets sharing the best ID according to step 1, for each group we trained the model on counts and labels from other groups to predict the second best ID of barcodes in the current group.
3. Doublets detection: to allow doublets detection to be specific and flexible, while accommodating genetic contributions ranging from balanced doublets to presence of cell and debris in the same drop, we implemented a method similar to the one adopted in ^22^. Starting from previous *m by g* contribution matrix, for every genotype *g* we define as negative droplets the ones that do not contain that genotype as best ID according to first step and fit a negative binomial distribution via fitdistrplus ^84^ R function on counts supporting private *g* alleles. We so used the 99% quantile of the fitted distribution as positivity threshold. Droplets positive to more than one ID are considered multiplets.
4. We finally took advantage of the mixed genotypes design to structure an added layer of a low quality droplets detection, to be used during consensus call aggregation.

We applied a Gaussian mixture model expectation maximisation algorithm (implemented through R MixTools ^85^ package) to separate droplets with ‘low’ and ‘high’ signal-to-noise by computing logFC between first and second best predicted IDs. We started by preparing a new contribution matrix similar to the one in passage 2, but considering only non-ambiguous loci between each possible pair of best and second best IDs in the dataset. Additionally, prior to logFC calculation we add pseudocounts, which mimics average ambient RNA contamination coming from each ID, calculated as average rate of reads deriving from other genotype’s private reads when they are not labelled as first nor second ID across all droplets (according to the contribution matrix again), similarly to the approach proposed in hashedDrops function from package MarioniLab/DropletUtils ^86,87^ this steps ensures that logFC is always defined for all droplets. Given the nature of the model, the resulting classification assumes the presence of two distinct populations that can be separated based on the proportion of the two IDs and given that it is computed after the doublets detection it will likely to detect those droplets that embed enough ambient RNA to pass cellranger emptyDrops filter, while it should not be used if any sort of prior filtering of low quality droplets has already been done.

### Aggregated call set up

#### Rationale and Tools

Given the widely diverse technical landscape deriving from samples’ multiplexing, particularly, but not-only bound to our specific *in vitro* system, we combined the output of multiple tools to retrieve the most accurate identity call and ensure maximum robustness across diverse technical conditions that could be challenging for the single tools. For this Consensus call, we combined 4 genetic demultiplexing tools: SCanSNP, Vireo, Souporcell and Demuxlet. Moreover we included scDblFinder ^43^ for non genetic doublets detection, and Dropkick ^44^, for low quality droplets detection. Software were embedded in a Snakamake pipeline ^88^, subsequently the results were aggregated into a weighted scoring system. All software provide a flexible output that allowed us to independently extract informations about overall assignment: i) overall assignment: droplet type (singlet, doublet, ambiguous, not assigned) plus best identity if the droplet was correctly assigned as singlet, ii) droplet type only, and iii) best guess of ID assuming singlet.

#### Metrics

*S*ince the endpoint of the aggregation is meant to be an identity confidently usable for downstream analyses we used as a general indicator for overall scoring the agreement between softwares’ best guess assuming singlet (ignoring at this stage the singlet / doublet prediction), this allowed us to maximise the sensitivity of tools while avoiding specific case/tools combinations that in some cases drives away the overall assignment. First, we assigned different weights to the software calls when computing the consensus score: +1 for Souporcell +1 for Vireo, +1.5 for Demuxlet +1.5 for SCanSNP. This choice was made to leverage the greater sensitivity of Demuxlet and SCanSNP best guesses assuming singlets even in hardest conditions (i.e. low quality drops, high amount of ambient-RNA (not shown). According to these weights the consensus score is computed for *g* Genotypes and *m* droplets, by summing the weighted score for *g* across tools. Next we set an “agreement threshold”, to be used downstream, as a measure of the stability of the assignment across tools. The “agreement” threshold for an ID was set to 3. E.g., same best ID predicted from SCanSNP and Demuxlet, or, same best ID predicted from Souporcell + Vireo + SCanSNP.

#### Low quality and Doublets

To ensure specific call for singlets and efficiently label doublets and low quality beads we proceeded through multi-tiered combined id assignment per barcode. At this stage we accounted for agreement meeting and re-introduced droplet-type information. First step was low quality droplet isolation: a droplet was labelled as “low quality” if a) it failed to reach agreement for any ID, and SCanSNP predicted it as low quality, or b) it failed to reach agreement for any ID, and dropkick predicted it as low quality. Subsequently we marked genetic doublets as droplets that a) were labelled as doublets by SCanSNP and by at least one more software or b) were labelled as doublet from all tools but SCanSNP and failed to meet agreement. Here we intentionally put SCanSNP on a different level with respect to other tools for the parsimony in calling doublets also in presence of challenging datasets with abundant ambient RNA. After the first two tiers the remaining cells classified as doublets from scDblFinder were considered transcriptomic doublets. At this point remaining barcodes meeting agreement were labelled as “safe singlets” and barcodes that did not were merged with previous “low quality”.

### *In silico* multiplexed bam creation

In order to explore the behaviour of the available demultiplexing tools we produced a set of *in silico* multiplexed BAM files starting from five individual SC RNA-seq datasets containing 1311, 8358, 10270, 3216, 2366 barcodes (filtered cellranger output) in the following manner: 1) Starting from filtered barcodes emitted by cellranger the overlapping ones across datasets were removed.

2A) Balanced in-silico datasets: maximum of 2000 barcodes per dataset were randomly selected among the unique ones to produce an overall balanced Barcode-ID list with 1311 barcodes for the first dataset and 2000 for the other ones sampling repeated 2 times). 2B) imbalanced datasets: only one of the 5 datasets was randomly sampled for 2000 barcodes, for the other ones all the non-overlapping barcodes were used, leading to the production of mostly imbalanced barcode-ID proportions (repeated 3 times changing the downsampled dataset). 3) At this point we obtained a sample-wise whitelist of barcodes to be sampled so obtain each of the five simulated dataset. We used the whitelists to filter each individual bam file and calculated the median reads, in case it exceeded the target median of 15000 it was subsampled accordingly using ‘samtools view -s .4’) Datasets were merged together using ‘gatk MergeSamFiles’. Starting from the 5 individual datasets we obtained 5 multiplexed (2 balanced and 3 imbalanced) BAM files. 5) In each multiplexed BAM, barcodes tag in aligned reads were swapped to reach 10% doublets portion. After the preparation of the *in silico* multiplexed datasets, we proceeded to the deconvolution using the available softwares. Since Souporcell does not use external reference for the demultiplexing the labels of predicted clusters were assigned using best matching ID from demuxlet. In the synthetic BAM we do not expect to see cross-genotypes ambient RNA background, that is used from SCanSNP to force mixture model fitting in order to separate low and high signal-to-noise containing droplets, therefore SCanSNP low quality detection was ignored.

### Power estimation for single cell eQTLs

We estimated the eQTL power using our R package scPower (v1.0.2) ^65^ for sample sizes between 25 and 200 and for number of cells per sample between 250 and 1,500, keeping the read depth as in our experiment. We fitted the required expression priors per cell type using the complete scRNA-seq dataset, combining the different multiplexing strategies and time points, and took effects from a previously published single cell eQTL study in IPS cells ^35^, excluding eQTLs from ROT treated cells. Genes were defined as expressed with at least 3 counts in at least 9.5% of the samples.

## Data and code availability

Full code used for the analyses can be retrieved at https://github.com/GiuseppeTestaLab/organoidMultiplexing_release. SCanSNP latest release and the docker link are available at https://github.com/GiuseppeTestaLab/SCanSNP.

---

**Fig. S1.**
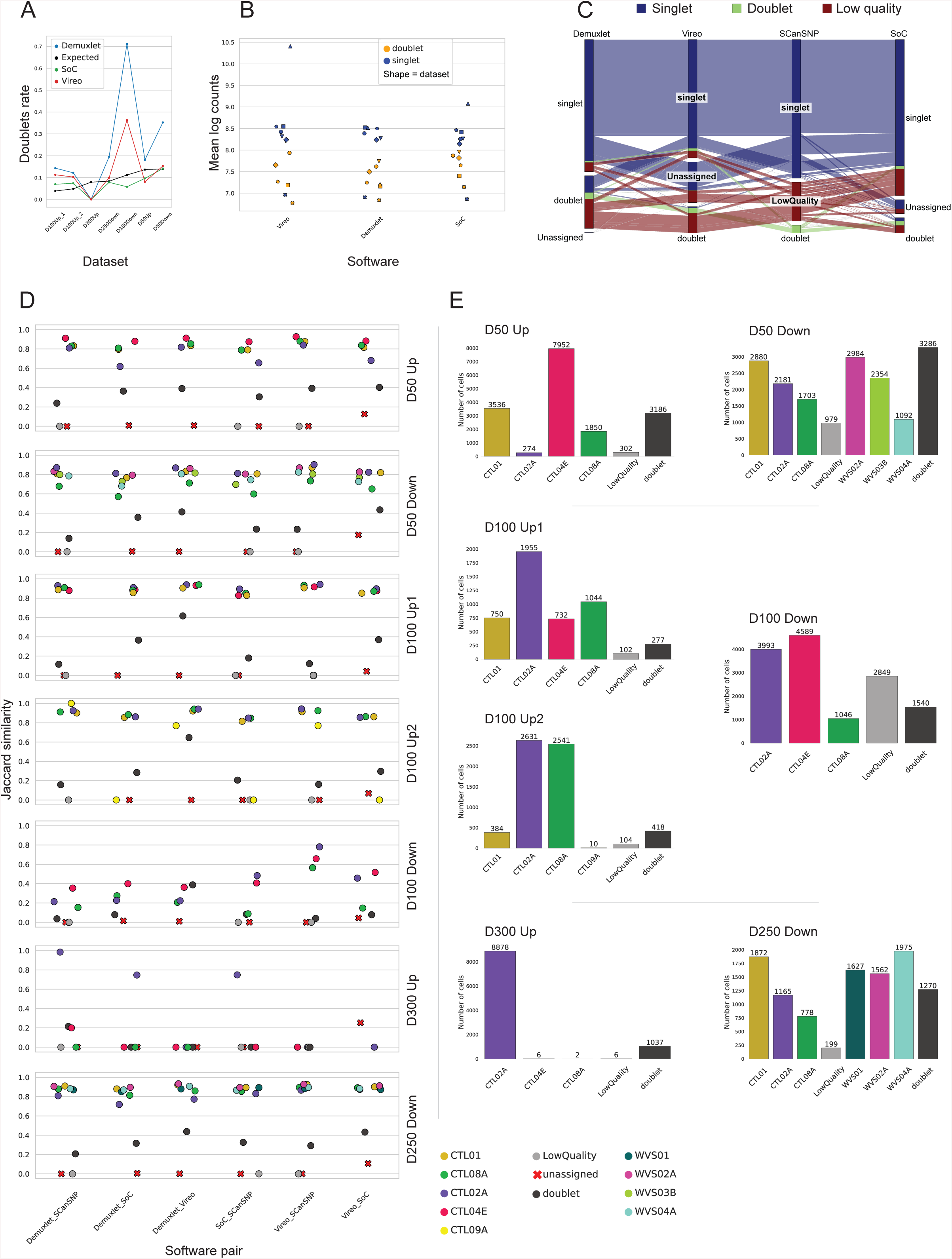
Demultiplexing performance assessment. **A)** Dublet rate by dataset and algorithm. Lines are coloured by demultiplexing algorithm, datasets (x axis) are ordered by number of retrieved cells. **B)** Average log counts distribution by demultiplexing algorithm. Dots are coloured by predicted singlet or doublet identity by each algorithm; the shape of the dots encodes each of the 7 datasets. **C)** Alluvial plot displaying Singlets, Doublets, Unassigned and Low quality classes mappings across demultiplexing algorithms. Cells (rows) are coloured according to SCanSNP assignment class. **D)** Pairwise agreement among software, divided by dataset. The agreement is expressed as a Jaccard similarity of each called identity between 2 software. **E)** Barplot representing number of cells by genotype according to the aggregated call prior to filtering. WVS01, WVS02A, WVS03B, WVS04A, CTL09A were not included in downstream analysis since there were no replicates across multiplexing modalities.

**Fig. S2.**
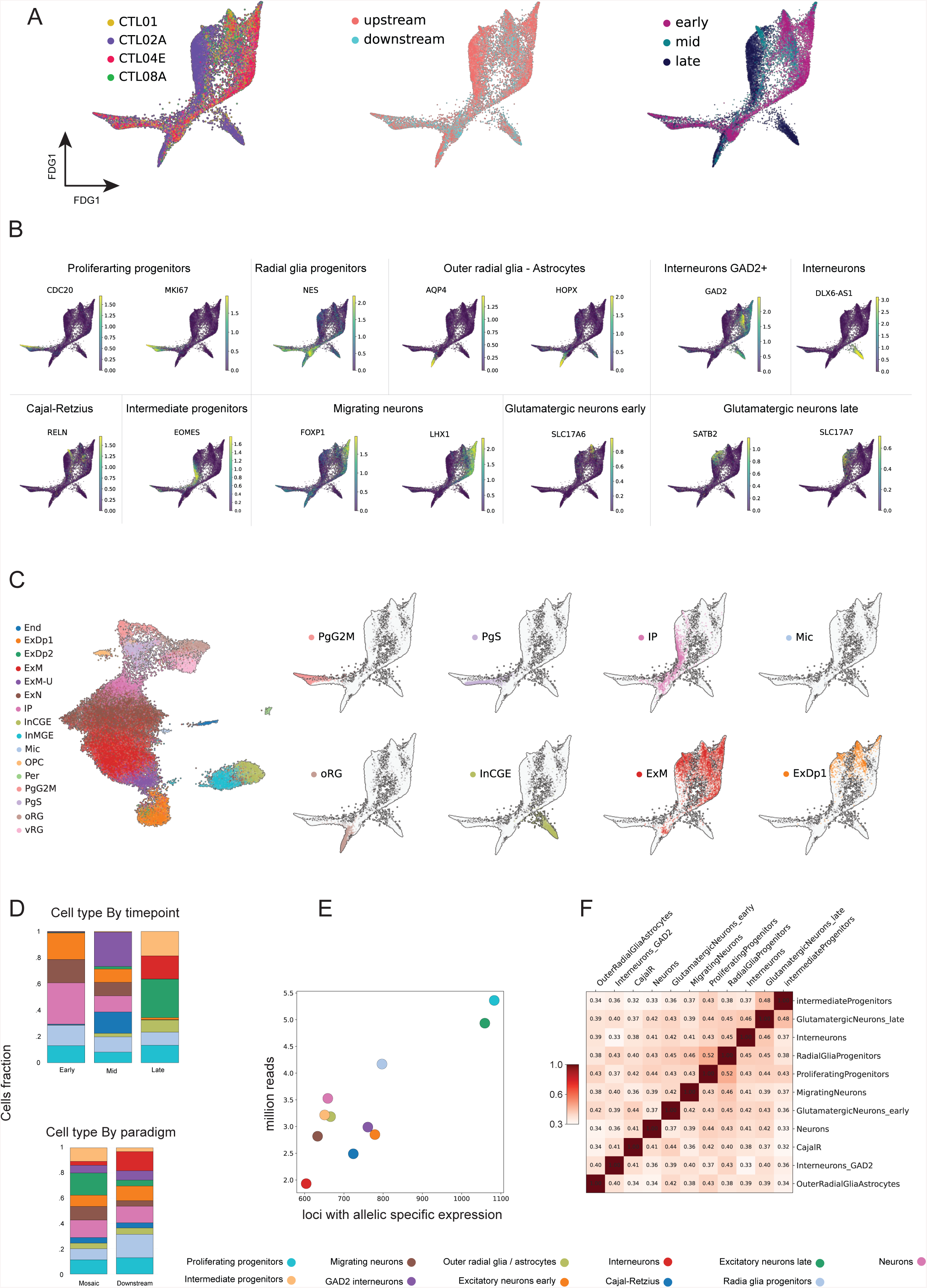
Single cell datasets characterization. **A)** Embedding of cells from all datasets of force-directed graph. From left to right cells are coloured by genotype, multiplexing paradigm and stage. **B)** Force-directed graph coloured by expression of relevant markers. Plotted markers are divided by the cell type they are most relevant for. **C)** Force-directed graph coloured by transferred label from *Poliudakis et al.* dataset. End: endothelial; ExDp1: soc excitatory deep-layer1; ExDp2: excitatory deep-layer2; vRG: ventral radial microglia; oRG: outer radial glia; ExN: newborn excitatory; ExM: maturing excitatory; ExM-U: e excitatory upper-layer-enriched; IP: intermediate progenitors; inCGE: interneurons caudal ganglionic eminence; inMGE: interneurons medial ganglionic eminence; Mic: microglia; OPC: oligodendrocyte precursors; Per: pericytes; PgG2M: G2M phase proliferating progenitors; PgGS: S phase proliferating progenitors; **D)** Plot of fraction of cells for each cell type, divided by timepoint (upper panel), and by multiplexing paradigm (lower panel). **E)** The scatterplot shows the number of loci with detected allele specific expression on the × axis by the total number of reads expressed in millions on the y axis, each dot represents a cell type. **F)** Spearman correlation on reads bringing alternative alleles / total reads (bValues) among the observed cell types. Correlation is calculated on loci that displayed allelic imbalance (binomial test fdr < 0.05) in at least one cell type and with at least 20 reads in each cell type.

**Fig. S3.**
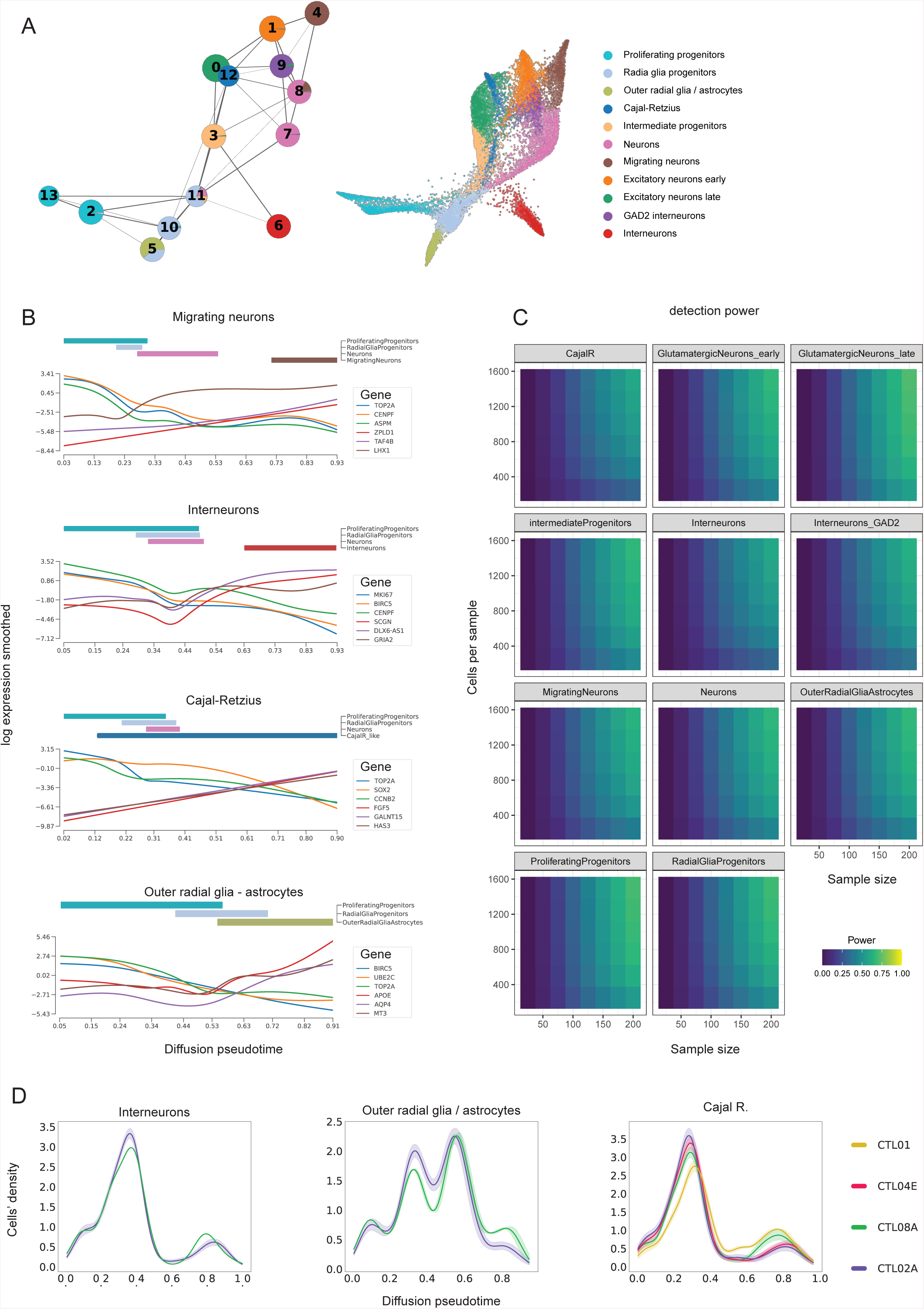
Developmental trajectory analysis and power analysis. **A)** On the left: Partition-based graph abstraction (PAGA) plot. Each circle represents a Paga cluster, circles are partitioned according to the fraction of cells per annotated celltype (shown as reference on the right side), weighted edges among Paga clusters encode their transcriptional similarity. **B)** Plot of smoothed gene expression - obtained via tradeseq - along pseudotime (methods). For each lineage the 3 most relevant decreasing and increasing genes (sorted by pVal and absolute logFC) are shown. Above each expression panel, bars coloured by cell type indicate the occupancy of each cell type along pseudotime. **C)** SCPOWER: Estimation of single cell eQTL power per specific cell type, depending on the sample size and numbers of cells per sample. **D)** Distribution of genotypes along pseudotime for Interneurons, Outer radial glia / astrocytes and Cajal R. lineages. Within each differentiation stage cells were balanced to the same amount across genotypes for unbiased comparison. If too few/no cells were retrieved at any differentiation stage the whole genotype was removed from the comparison. Faded colour shows 1 standard deviation across random subsampling iterations

**Fig. S4.**
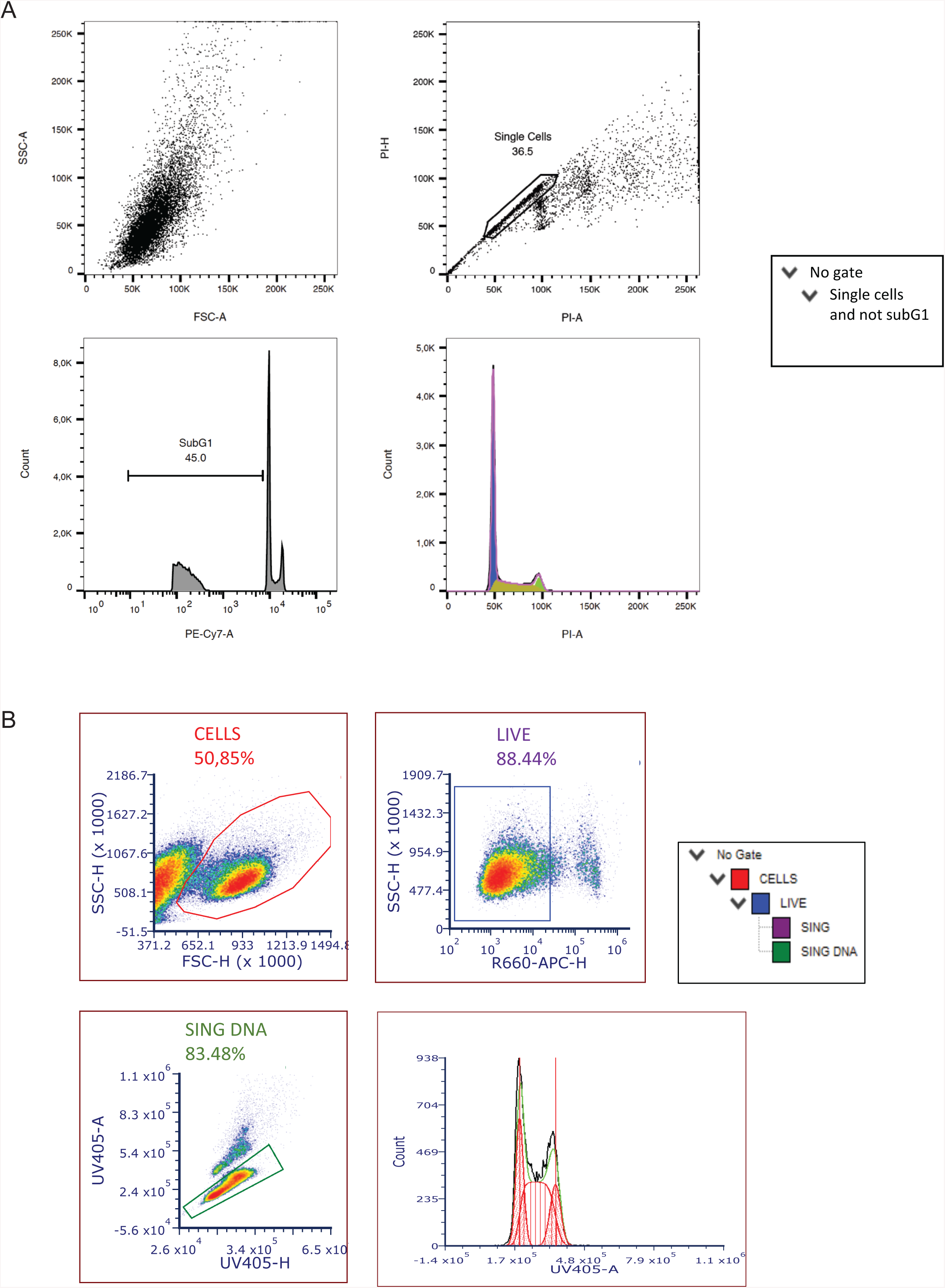
Cell cycle analysis gating strategy. **A)** Replicate 1: gating strategy for longitudinal cell cycle analysis of ethanol fixed, Propidium Iodide stained dissociated organoids. **B)** Replicate 2: gating strategy for longitudinal cell cycle analysis of freshly dissociated organoids stained with Hoecst33342 (total DNA content) and ToPro dye (live/dead).

**Table S1. Longitudinal cell cycle analysis of pure lines and mosaic organoids**

Percentages of cells in each phase of the cell cycle (G0/G1, S, G2/M) at multiple differentiation timepoints.

**Table S2. Annotation markers an Cluster markers**

Main markers used in the annotation phase. Top DEGs among annotated clusters (Wilcoxon rank sum test - one vs all - benjamini-hochberg < 0.05). Additional HVGs to be considered in the PCA..

**Table S3. Developmental trajectories transcriptional dynamics**

Most relevant, dynamically-changing genes along pseudotime (methods). Each sheet reports a developmental branch.

**Table S4. Cells density per genotype along pseudotime**

Cells density along pseudotime per genotype. Mean and standard deviations are reported for 50 downsampling iterations. Each sheet reports a developmental branch

**Table S5. SCeQTLs in migrating neurons**

List of SCeQTLs in Jerber et. al found in migrating neurons lineage between the two genotype groups (CTL01 & CTL02A vs CTL04E & CTL08A)

**Table S6. hiPSC lines employed**

Nomenclature of the hiPSC lines employed and their corresponding nomenclature in the human pluripotent stem cell registry (hPSCreg) ^89^, origin and sex of the lines.

CTL08A, CTL01, CTL02A, CTL04E were included in the experimental design for the single cell analysis of neurodevelopmental cell populations and trajectories. CTL09A was originally included in only one experiment to generate mosaic organoids but not retrieved at D100 profiling and was hence not included in the single cell datasets analysed across replicates for all timepoints. WVS01H, WVS02A, WVS03B, WVS04A were not part of the study design aimed at evaluating upstream versus downstream multiplexing for the detection of neurodevelopmental trajectories. They are listed here only insofar as they were included, for maximising sequencing cost-effectiveness, in the preparation of the downstream multiplexing libraries and hence harnessed as additional lines solely to benchmark the demultiplexing pipelines.

## References

1. Silbereis, J. C., Pochareddy, S., Zhu, Y., Li, M. & Sestan, N. The Cellular and Molecular Landscapes of the Developing Human Central Nervous System. Neuron 89, 248–268 (2016).

2. Cheroni, C., Caporale, N. & Testa, G. Autism spectrum disorder at the crossroad between genes and environment: contributions, convergences, and interactions in ASD developmental pathophysiology. Mol. Autism 11, 69 (2020).

3. Tărlungeanu, D. C. & Novarino, G. Genomics in neurodevelopmental disorders: an avenue to personalized medicine. Exp. Mol. Med. 50, 1–7 (2018).

4. Hyman, S. E. The daunting polygenicity of mental illness: making a new map. Philos. Trans. R. Soc. Lond. B Biol. Sci. 373, (2018).

5. López-Tobón, A. et al. Human Cortical Organoids Expose a Differential Function of GSK3 on Cortical Neurogenesis. Stem Cell Reports 13, 847–861 (2019).

6. Lopez-Tobon, A. et al. GTF2I dosage regulates neuronal differentiation and social behavior in 7q11.23 neurodevelopmental disorders. bioRxiv 2022.10.10.511434 (2022) doi:10.1101/2022.10.10.511434.

7. Mihailovich, M. et al. 7q11.23 CNV alters protein synthesis and REST-mediated neuronal intrinsic excitability. bioRxiv 2022.10.10.511483 (2022) doi:10.1101/2022.10.10.511483.

8. Marangon, D. et al. Novel in vitro Experimental Approaches to Study Myelination and Remyelination in the Central Nervous System. Front. Cell. Neurosci. 15, 748849 (2021).

9. Drakulic, D. et al. Copy number variants (CNVs): a powerful tool for iPSC-based modelling of ASD. Mol. Autism 11, 42 (2020).

10. Villa, C. E. et al. CHD8 haploinsufficiency links autism to transient alterations in excitatory and inhibitory trajectories. Cell Rep. 39, 110615 (2022).

11. Caporale, N. et al. From cohorts to molecules: Adverse impacts of endocrine disrupting mixtures. Science 375, eabe8244 (2022).

12. Corsini, N. S. & Knoblich, J. A. Human organoids: New strategies and methods for analyzing human development and disease. Cell 185, 2756–2769 (2022).

13. Kelley, K. W. & Pașca, S. P. Human brain organogenesis: Toward a cellular understanding of development and disease. Cell 185, 42–61 (2022).

14. Eichmüller, O. L. & Knoblich, J. A. Human cerebral organoids - a new tool for clinical neurology research. Nat. Rev. Neurol. 18, 661–680 (2022).

15. Cheroni, C. et al. Benchmarking brain organoid recapitulation of fetal corticogenesis. Transl. Psychiatry 12, 520 (2022).

16. Koi, P. Genetics on the neurodiversity spectrum: Genetic, phenotypic and endophenotypic continua in autism and ADHD. Stud. Hist. Philos. Sci. 89, 52–62 (2021).

17. Baron-Cohen, S. Editorial Perspective: Neurodiversity - a revolutionary concept for autism and psychiatry. J. Child Psychol. Psychiatry 58, 744–747 (2017).

18. Pretzsch, C. M. et al. Neurobiological Correlates of Change in Adaptive Behavior in Autism. Am. J. Psychiatry 179, 336–349 (2022).

19. Rajewsky, N. et al. LifeTime and improving European healthcare through cell-based interceptive medicine. Nature 587, 377–386 (2020).

20. Cuomo, A. S. E., Nathan, A., Raychaudhuri, S., MacArthur, D. G. & Powell, J. E. Single-cell genomics meets human genetics. Nat. Rev. Genet. 24, 535–549 (2023).

21. De Donno, C. et al. Population-level integration of single-cell datasets enables multi-scale analysis across samples. bioRxiv 2022.11.28.517803 (2022) doi:10.1101/2022.11.28.517803.

22. Stoeckius, M. et al. Cell Hashing with barcoded antibodies enables multiplexing and doublet detection for single cell genomics. Genome Biol. 19, 224 (2018).

23. Stoeckius, M. et al. Simultaneous epitope and transcriptome measurement in single cells. Nat. Methods 14, 865–868 (2017).

24. Gehring, J., Hwee Park, J., Chen, S., Thomson, M. & Pachter, L. Highly multiplexed single-cell RNA-seq by DNA oligonucleotide tagging of cellular proteins. Nat. Biotechnol. 38, 35–38 (2020).

25. Hwang, B. et al. SCITO-seq: single-cell combinatorial indexed cytometry sequencing. Nat. Methods 18, 903–911 (2021).

26. Datlinger, P. et al. Ultra-high-throughput single-cell RNA sequencing and perturbation screening with combinatorial fluidic indexing. Nat. Methods 18, 635–642 (2021).

27. Kang, H. M. et al. Multiplexed droplet single-cell RNA-sequencing using natural genetic variation. Nat. Biotechnol. 36, 89–94 (2018).

28. Heaton, H. et al. Souporcell: robust clustering of single-cell RNA-seq data by genotype without reference genotypes. Nat. Methods 17, 615–620 (2020).

29. Vireo: Bayesian demultiplexing of pooled single-cell RNA-seq data without genotype reference - Genome Biology. BioMed Central https://genomebiology.biomedcentral.com/articles/10.1186/s13059-019-1865-2,.

30. Weber, L. M. et al. Genetic demultiplexing of pooled single-cell RNA-sequencing samples in cancer facilitates effective experimental design. Gigascience 10, (2021).

31. Yazar, S. et al. Single-cell eQTL mapping identifies cell type-specific genetic control of autoimmune disease. Science 376, eabf3041 (2022).

32. Perez, R. K. et al. Single-cell RNA-seq reveals cell type-specific molecular and genetic associations to lupus. Science 376, eabf1970 (2022).

33. Srivatsan, S. R. et al. Massively multiplex chemical transcriptomics at single-cell resolution. Science 367, 45–51 (2020).

34. Cuomo, A. S. E. et al. Single-cell RNA-sequencing of differentiating iPS cells reveals dynamic genetic effects on gene expression. Nat. Commun. 11, 810 (2020).

35. Jerber, J. et al. Population-scale single-cell RNA-seq profiling across dopaminergic neuron differentiation. Nat. Genet. 53, 304–312 (2021).

36. Mitchell, J. M. et al. Mapping genetic effects on cellular phenotypes with ‘cell villages’. bioRxiv 2020.06.29.174383 (2020) doi:10.1101/2020.06.29.174383.

37. Wells, M. F. et al. Natural variation in gene expression and viral susceptibility revealed by neural progenitor cell villages. Cell Stem Cell 30, 312–332.e13 (2023).

38. Neavin, D. R. et al. A village in a dish model system for population-scale hiPSC studies. Nat. Commun. 14, 3240 (2023).

39. Pașca, S. P., et al. A nomenclature consensus for nervous system organoids and assembloids. Nature 609, 907–910 (2022).

40. Birey, F. et al. Assembly of functionally integrated human forebrain spheroids. Nature 545, 54–59 (2017).

41. Bizzotto, S. et al. Landmarks of human embryonic development inscribed in somatic mutations. Science 371, 1249–1253 (2021).

42. Huang, Y., McCarthy, D. J. & Stegle, O. Vireo: Bayesian demultiplexing of pooled single-cell RNA-seq data without genotype reference. Genome Biol. 20, 273 (2019).

43. Germain, P.-L., Lun, A., Macnair, W. & Robinson, M. D. Doublet identification in single-cell sequencing data using scDblFinder. F1000Res. 10, 979 (2021).

44. Heiser, C. N., Wang, V. M., Chen, B., Hughey, J. J. & Lau, K. S. Automated quality control and cell identification of droplet-based single-cell data using dropkick. Genome Res. 31, 1742–1752 (2021).

45. Polioudakis, D. et al. A Single-Cell Transcriptomic Atlas of Human Neocortical Development during Mid-gestation. Neuron 103, 785–801.e8 (2019).

46. Tanaka, Y., Cakir, B., Xiang, Y., Sullivan, G. J. & Park, I.-H. Synthetic Analyses of Single-Cell Transcriptomes from Multiple Brain Organoids and Fetal Brain. Cell Rep. 30, 1682–1689.e3 (2020).

47. Uzquiano, A. et al. Proper acquisition of cell class identity in organoids allows definition of fate specification programs of the human cerebral cortex. Cell 185, 3770–3788.e27 (2022).

48. Jourdon, A. et al. Modeling idiopathic autism in forebrain organoids reveals an imbalance of excitatory cortical neuron subtypes during early neurogenesis. Nat. Neurosci. (2023) doi:10.1038/s41593-023-01399-0.

49. Dann, E., Henderson, N. C., Teichmann, S. A., Morgan, M. D. & Marioni, J. C. Differential abundance testing on single-cell data using k-nearest neighbor graphs. Nat. Biotechnol. 40, 245–253 (2022).

50. Cleary, S. & Seoighe, C. Perspectives on Allele-Specific Expression. Annu Rev Biomed Data Sci 4, 101–122 (2021).

51. Zhao, D., Lin, M., Pedrosa, E., Lachman, H. M. & Zheng, D. Characteristics of allelic gene expression in human brain cells from single-cell RNA-seq data analysis. BMC Genomics 18, 860 (2017).

52. Tritschler, S. et al. Concepts and limitations for learning developmental trajectories from single cell genomics. Development 146, (2019).

53. Deconinck, L., Cannoodt, R., Saelens, W., Deplancke, B. & Saeys, Y. Recent advances in trajectory inference from single-cell omics data. Current Opinion in Systems Biology 27, 100344 (2021).

54. Wolf, F. A. et al. PAGA: graph abstraction reconciles clustering with trajectory inference through a topology preserving map of single cells. Genome Biol. 20, 59 (2019).

55. Haghverdi, L., Büttner, M., Wolf, F. A., Buettner, F. & Theis, F. J. Diffusion pseudotime robustly reconstructs lineage branching. Nat. Methods 13, 845–848 (2016).

56. Van den Berge, K., et al. Trajectory-based differential expression analysis for single-cell sequencing data. Nat. Commun. 11, 1201 (2020).

57. Pebworth, M.-P., Ross, J., Andrews, M., Bhaduri, A. & Kriegstein, A. R. Human intermediate progenitor diversity during cortical development. Proc. Natl. Acad. Sci. U. S. A. 118, (2021).

58. Braun, E. et al. Comprehensive cell atlas of the first-trimester developing human brain. bioRxiv 2022.10.24.513487 (2022) doi:10.1101/2022.10.24.513487.

59. Kriegstein, A., Noctor, S. & Martínez-Cerdeño, V. Patterns of neural stem and progenitor cell division may underlie evolutionary cortical expansion. Nat. Rev. Neurosci. 7, 883–890 (2006).

60. Sollis, E. et al. The NHGRI-EBI GWAS Catalog: knowledgebase and deposition resource. Nucleic Acids Res. 51, D977–D985 (2023).

61. Aguet, F. et al. Molecular quantitative trait loci. Nature Reviews Methods Primers 3, 1–22 (2023).

62. van der Wijst, M. et al. The single-cell eQTLGen consortium. Elife 9, (2020).

63. GTEx Consortium et al. Genetic effects on gene expression across human tissues. Nature 550, 204–213 (2017).

64. Cuomo, A. S. E. et al. CellRegMap: a statistical framework for mapping context-specific regulatory variants using scRNA-seq. Mol. Syst. Biol. 18, e10663 (2022).

65. Schmid, K. T. et al. scPower accelerates and optimizes the design of multi-sample single cell transcriptomic studies. Nat. Commun. 12, 6625 (2021).

66. Chapelle, J. et al. Dissecting the Shared and Context-Dependent Pathways Mediated by the p140Cap Adaptor Protein in Cancer and in Neurons. Front Cell Dev Biol 7, 222 (2019).

67. Grasso, S. et al. The SRCIN1/p140Cap adaptor protein negatively regulates the aggressiveness of neuroblastoma. Cell Death Differ. 27, 790–807 (2020).

68. Fleck, J. S. et al. Inferring and perturbing cell fate regulomes in human brain organoids. Nature (2022) doi:10.1038/s41586-022-05279-8.

69. Esk, C. et al. A human tissue screen identifies a regulator of ER secretion as a brain-size determinant. Science 370, 935–941 (2020).

70. CRISPR screens in 3D assembloids reveal disease genes associated with human interneuron development.

71. Single-cell brain organoid screening identifies developmental defects in autism.

72. Elorriaga, V., Pierani, A. & Causeret, F. Cajal-retzius cells: Recent advances in identity and function. Curr. Opin. Neurobiol. 79, 102686 (2023).

73. Sloan, S. A., Andersen, J., Pașca, A. M., Birey, F. & Pașca, S. P. Generation and assembly of human brain region-specific three-dimensional cultures. Nat. Protoc. 13, 2062–2085 (2018).

74. Delgado, R. N. et al. Individual human cortical progenitors can produce excitatory and inhibitory neurons. Nature 601, 397–403 (2022).

75. Seah, C. et al. Modeling gene × environment interactions in PTSD using human neurons reveals diagnosis-specific glucocorticoid-induced gene expression. Nat. Neurosci. 25, 1434–1445 (2022).

76. Yoon, S.-J. et al. Reliability of human cortical organoid generation. Nat. Methods 16, 75–78 (2019).

77. Dobin, A. et al. STAR: ultrafast universal RNA-seq aligner. Bioinformatics 29, 15–21 (2013).

78. Poplin, R., et al. Scaling accurate genetic variant discovery to tens of thousands of samples. bioRxiv (2017) doi:10.1101/201178.

79. Wolf, F. A., Angerer, P. & Theis, F. J. SCANPY: large-scale single-cell gene expression data analysis. Genome Biol. 19, 15 (2018).

80. Korsunsky, I. et al. Fast, sensitive and accurate integration of single-cell data with Harmony. Nat. Methods 16, 1289–1296 (2019).

81. Proceedings of the python in science conference (SciPy): Exploring network structure, dynamics, and function using NetworkX. https://conference.scipy.org/proceedings/SciPy2008/paper_2/.

82. Haghverdi, L., Buettner, F. & Theis, F. J. Diffusion maps for high-dimensional single-cell analysis of differentiation data. Bioinformatics 31, 2989–2998 (2015).

83. Virtanen, P. et al. SciPy 1.0: fundamental algorithms for scientific computing in Python. Nat. Methods 17, 261–272 (2020).

84. Delignette-Muller, M. L. & Dutang, C. fitdistrplus: An R Package for Fitting Distributions. J. Stat. Softw. 64, 1–34 (2015).

85. Benaglia, T., Chauveau, D., Hunter, D. R. & Young, D. S. mixtools: An R Package for Analyzing Mixture Models. J. Stat. Softw. 32, 1–29 (2010).

86. Lun, A. T. L. et al. EmptyDrops: distinguishing cells from empty droplets in droplet-based single-cell RNA sequencing data. Genome Biol. 20, 63 (2019).

87. Griffiths, J. A., Richard, A. C., Bach, K., Lun, A. T. L. & Marioni, J. C. Detection and removal of barcode swapping in single-cell RNA-seq data. Nat. Commun. 9, 2667 (2018).

88. Mölder, F. et al. Sustainable data analysis with Snakemake. F1000Res. 10, 33 (2021).

89. Seltmann, S. et al. hPSCreg--the human pluripotent stem cell registry. Nucleic Acids Res. 44, D757–63 (2016).

